# Genetic Dependence and Genetic Diseases

**DOI:** 10.1101/2023.08.02.551736

**Authors:** Bin Li, Wen-Jun Bian, Peng Zhou, Jie Wang, Cui-Xia Fan, Hai-Qing Xu, Lu Yu, Na He, Yi-Wu Shi, Tao Su, Yong-Hong Yi, Wei-Ping Liao

**Author notes:** Correspondence: Prof. Wei-Ping Liao (,), Department of Neurology, Institute of Neuroscience, Key Laboratory of Neurogenetics and Channelopathies of Guangdong Province and the Ministry of Education of China, the Second Affiliated Hospital, Guangzhou Medical University, Guangzhou, 510260, China. These authors contributed equally to the manuscript (Bin Li, Wen-Jun Bian, Peng Zhou).

## Abstract

The human life depends on the function of proteins that are encoded by about twenty-thousand genes. The gene-disease associations in majority genes are unknown and the mechanisms underlying pathogenicity of genes/variants and common diseases remain unclear. We studied how human life depends on the genes, i.e., the genetic-dependence, which was classified as genetic-dependent nature (GDN, vital consequence of abolishing a gene), genetic-dependent quantity (GDQ, quantitative genetic function required for normal life), and genetic-dependent stage (GDS, temporal expression pattern). Each gene differs in genetic-dependent features, which determines the gene-disease association extensively. The GDN is associated with the pathogenic potential/feature of genes and the strength of pathogenicity. The GDQ-damage relation determines the pathogenicity of variants and subsequently the pathogenic genotype, phenotype spectrum, and inheritance of variants. The GDS is mainly associated with the onset age/evolution/outcome and the nature of genetic disorders (disease/susceptibility). The varied and quantitative genetic-dependent feature of genome explains common mild phenotype/susceptibility. The genetic-dependence discloses the mechanisms underlying pathogenicity of gene/variants and common diseases.

**One sentence summary:** Genetic dependent feature differs in genes and determines pathogenicity of genes/variants and the clinical features of genetic diseases.

## Introduction

The normal function of proteins, which are encoded by about twenty thousand genes ^1^, is the basis of human life. Variations in genes not only make us unique but also are associated with diseases. However, it is unknown to what extent genetic variations are linked with personal health, which is the fourth of the 125 big questions facing science ^2^ and also a substantial challenge of medicine. Based on classical Mendelian genetics, 4,620 genes have been reported to be potentially associated with human diseases (https://omim.org/), mostly rare diseases. The genetic mechanisms underlying common diseases and the determinants underlying the pathogenicity of genes is unknown. On the other hand, though a large number of sequence variants can be detected in each individual by next generation sequencing, determination of the pathogenicity of the variants remains a major challenge ^3^. Variants with the same damaging effect, such as nonsense variants that lead to haploinsufficiency, may be pathogenic for some genes but non-pathogenic for others. Apparently, factors other than the damage effect contribute to the pathogenicity of variants.

Normal life depends on the function of genes, which is conceptualized as “genetic dependence”. In a general sense, genetic dependence is defined as how normal life depends on the function of a gene. To disclose the relationship between genetic variations and human health, we studied how normal life depends on the function of each gene. The genetic-dependent features were broadly analyzed and classified as genetic dependent nature (GDN), genetic dependent quantity (GDQ), and genetic dependent stage (GDS). The results show that each gene differs in genetic dependent feature that determines the gene-disease association extensively, including the pathogenic potential and pathogenic feature of genes, the pathogenicity of genetic variants, and several critical clinical characteristics of the genetic diseases, such as onset age, evolution, and nature of genetic disorders (disease/susceptibility). The quantitative nature of genetic dependence, together with the phenotype nature and spectrum determined by GDN and GDS, explains the occurrence of common diseases. This study suggested the genetic dependence as a principle of genetics that potentially discloses the genetic mechanisms underlying the links between genetic variations and human health, providing the bases of the individualized medicine in the future.

## Materials and Methods

### Definitions and data analysis

We studied the 19711 protein-coding genes in human genome (Hugo Gene Nomenclature Committee, www.genenames.org). The collected and analyzed information included data on expression, genetic knockouts (KOs), genetic diseases, bioinformatics, and related experiments or functional studies. We systematically studied the genetic-dependent features of protein-coding genes and their relationships with human genetic diseases.

### GDN

Each gene has a distinct role. Therefore, the absence of the genes has impact of different natures on the normal life. Based on the consequences of abolishing the genes (gene knock-out, KO), the GDN of genes can be classified as vital, obligatory, fractional, or dispensable/replaceable, where 1) vital: abolishment of the gene leads to embryonic or early death; 2) obligatory: abolishment of the gene causes abnormal phenotypes (diseases); 3) fractional: abolishment of the gene causes detectable mild alterations, including physiological, biochemical, cellular, and organizational abnormalities, but not abnormal physical phenotypes; 4) dispensable/replaceable: abolishment of the gene has no detectable effect.

In this study, evaluation of the GDN was based mainly on evidence from KO mice, which are strikingly similar to humans at the genomic level and in regards to pathophysiological aspects of diseases ^4^. Information about genetic KOs was retrieved from Mouse Genome Informatics (MGI, http://www.informatics.jax.org/). Data from targeted KO was analyzed and positive findings were used for classification of GDQ. We also searched for KO information from public literature, including data on mice and other animal species, such as zebrafish and drosophila.

### GDQ and quantitative genetic-dependent range (GDR)

The human genome is diploid. Factitiously, the full scale of the functional potential of a gene, which is contributed by two copies of the gene, was quantified as 100%. Normally, only partial functioning of the two copies of a gene is required to maintain its biophysiological function, while a considerable portion of it serves as reserve. The lowest limit of the required quantity of genetic function was defined as the GDQ. When genetic damaging effect exceeds the GDQ, phenotype occurs. The GDQ was therefore determined by analyzing the correlation between the level of functional deficiency and the occurrence of phenotype, such as the phenotype of gene KO and the inheritance pattern of genetic diseases.

From the perspective of KO, the GDQ was classified as >50% and <50%, where GDQ >50% was indicated when heterozygous KO caused an abnormal phenotype; GDQ <50% was indicated when heterozygous KO was asymptomatic whereas homozygous KO resulted in abnormal phenotypes or embryonic/early death.

For genes that were associated with genetic diseases caused by loss of function (LOF) mechanism, the GDQ was grossly inferred by the inheritance pattern. In dominant genetic disorders, impairment of one of the two copies of the gene results in disease, indicating that the GDQ of the gene is >50%; whereas in recessive disorders, bi-allelic variants are required to cause disease, indicating the GDQ is <50%.

However, the GDQs of genes in genome would be a continuous distribution and include genes with GDQs of ∼50%, which was suggested when: a) monoallelic variants with severe damage (haploinsufficiency) were pathogenic but were associated with mild phenotypes, whereas bi-allelic variants with relatively less damage were potentially pathogenic or were associated with severer phenotypes (higher than but close to 50%); and b) besides the bi-allelic variants, monoallelic variants with LOF and dominant negative effect (or similar additional effect) resulted in diseases (lower than but close to 50%).

More precise GDQs were determined based on quantitative correlations between damage effect of variants and occurrence of phenotype, which were obtained from studies on residual activity, mosaicism (the mutation load) ^5^, and splicing analysis (normal/abnormal mRNA transcript ratios) ^6^.

After exceeding the GDQ, the functional damage may quantitatively correlate with the phenotype severity within a range, which was defined as the genetic-dependent range (GDR) of the gene.

Assessment of the GDQ and GDR was based on comprehensive analysis of published literatures from OMIM (www.omim.org), HGMD (www.hgmd.cf.ac.uk), and PubMed (www.pubmed.ncbi.nlm.nih.gov).

### GDS

Genes may be expressed and function throughout the life span, or in a specific period of developmental stage, during which the life depends on. In this study, the GDS was inferred from the temporal expression of genes at different development stages. Information on gene expression was retrieved from UniGene (https://www.ncbi.nlm.nih.gov/UniGene). The developmental stages were classified as: 1) prenatal, including the embryoid body, blastocyst, and fetal stages; 2) early life, including the neonatal and infant stages; and 3) mature, including the juvenile to adult stages. The expression levels of different stages were semi-quantified and ranked from “0” to “3”, with the highest level being “3” and undetectable being “0”. Expression levels within 80% to 100% of the highest value were qualified as “3”; expression levels of 50% to 80% of the highest value were defined as “2”; and those of less than 50% of the highest level and more than 0 were quantified as “1”.

We analyzed and presented the gene expression levels across developmental stages, e.g., 3-1-3/2 indicates the levels at the prenatal, early life, juvenile/adult stages. The GDSs were classified into five patterns, i.e., 1) the whole life stable, 2) predominantly mature (increasing-shape), 3) predominantly prenatal (decreasing-shape), 4) early life (“A”-shape), and 5) predominantly prenatal and mature (“V”-shape) patterns.

### The database

A web interface of GD&P (www.gdap.org.cn) was developed to present information on the genetic-dependent features and pathogenic features of each gene, as well as relevant data required to evaluate the pathogenicity of variants.

### Statistical analysis

Statistical analysis was performed with the SPSS version 20.0. The Kruskal-Wallis test was used to estimate the significance of correlations. Chi-square analysis was applied to compare the difference of frequencies. Values of p <0.05 (two-sided) were considered significant.

## Results

### The GDN and pathogenic potential/feature of genes

Among the 10,989 genes with KO information available, the genes with GDN of vital, obligatory, fractional, and dispensable/replaceable were 4,455 (40.5%), 4,889 (44.5%), 1,205 (11.0%), and 440 (4.0%), respectively (Figure 1a). We studied the role of the GDN in gene-disease associations. Among the genes associated with monogenic diseases in the OMIM (www.omim.org, until April 2022), the overwhelming majority were of vital (58.5%) and obligatory (37.7%) in GDN, while only 3.0% genes were fractional and 0.8% genes were dispensable/replaceable (Figure 1b, left panel), suggesting that the GDN is closely related to the pathogenic potential of genes, and genes with GDN of vital were mostly frequent in the genes with monogenic diseases of human.

**Figure 1.**
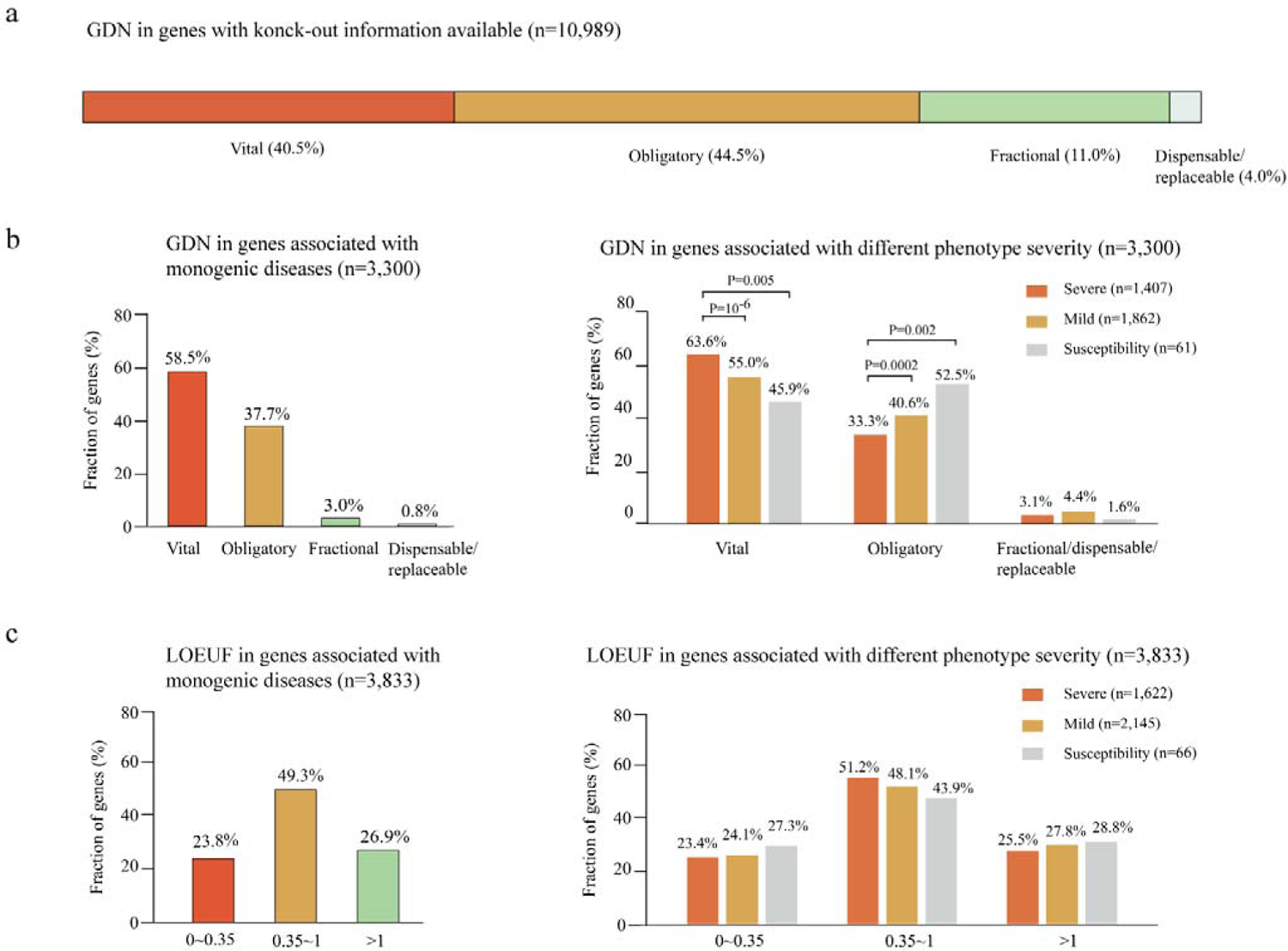
The genetic dependent nature (GDN) and its relationship with the pathogenic potential of genes. **(a)** GDNs in genes with knock-out information available (n=10989, including 10,950 from MGI and 39 from other sources). **(b)** GDNs in genes associated with monogenic diseases. These included phenotype-associated genes obtained from the OMIM database, with exclusion of genes associated with 1) non-disease phenotypes, 2) phenotypes associated with single-nucleotide polymorphisms, 3) polygenic diseases, 4) diseases associated with somatic mutations, and 5) phenotypic modifier roles. The overwhelming majority of the disease-associated genes (>95%) were of vital and obligatory in GDN (left). GDNs in genes associated with phenotype severity (right). Genes with GDN of vital presented a significantly higher proportion in severe phenotypes; and genes with GDN of obligatory had a significantly higher proportion in susceptibility and mild phenotypes. Chi-square statistical analysis was applied to compare the difference of frequencies. **(c)** Loss of function (LOF) observed/expected upper bound fraction index (LOEUF) in genes associated with monogenic diseases. Genes with LOEUF <0.35 were predicted to be highly intolerant to LOF, whereas LOEUF >1 were tolerant to LOF. Only a small proportion of the disease-associated genes (23.8%) were of <0.35 in LOEUF (left). There was no significant correlation between LOEUF value and phenotype severity (right).

We then analyzed the correlation between the GDN and phenotype severity, which is potentially associated with the strength of pathogenicity of the genes. The disease phenotypes were classified into severe, mild, and susceptibility, according to the nature and impact to human health of the diseases. The severe phenotypes were those with lethal effect or described to have severe impact on human health; and the others were classified as mild phenotype. The susceptibility genes were those defined in the OMIM (www.omim.org). Genes with GDN of vital accounted a significantly higher proportion in severe phenotypes (63.6%) than that in mild phenotypes (55.0%) or phenotypes of susceptibility (45.9%). In contrast, genes with GDN of obligatory had a significantly higher proportion in susceptibility (52.5%) and mild (40.6%) phenotypes than that in severe phenotypes (33.3%) (Figure 1b, right panel).

From the perspective of the human genome, the genes with GDN of obligatory were mostly common (44.5%). Complete functional abolishment of such genes would result in abnormal phenotypes, instead of lethality; whereas partial impairment of the genes, such as haploinsufficiency, would be associated with further milder phenotype or susceptibility alteration. Therefore, the pathogenicity of genes with GDN of obligatory would potentially be moderate, typically as *CACNA1H* and *EFHC1* that are associated with common mild generalized epilepsies but in long-standing dispute in pathogenicity ^7^. However, these genes are potentially more relevant as sources of common diseases ^8^.

We further analyzed the pathogenic mechanisms of the genes with fractional and dispensable/replaceable GDNs (Table S1 and Table 1). There were 64 genes of fractional GDN with confirmed pathogenicity (www.gdap.org.cn) ^9^, among which 32 genes were associated with pathogenic mechanisms other than loss of function (LOF), including toxic effects, gain of function (GOF), LOF with dominant negative effects, and specific pathogenic mechanism (*PCDH19* with “heterogeneity in cells”) ^10^. The consistently mild and later onset phenotypes explained 18 and 9 genes, respectively (Table S1). In only 5 genes of fractional GDN, the pathogenic potential was unexplainable. Similarly, pathogenicity of the 11 genes of dispensable/replaceable GDN were explained by pathogenic mechanisms other than LOF (6 genes), the consistently mild/late-onset phenotypes (4 genes), and the genome difference between human and mouse (2 genes) (Table 1).

**Table 1.**
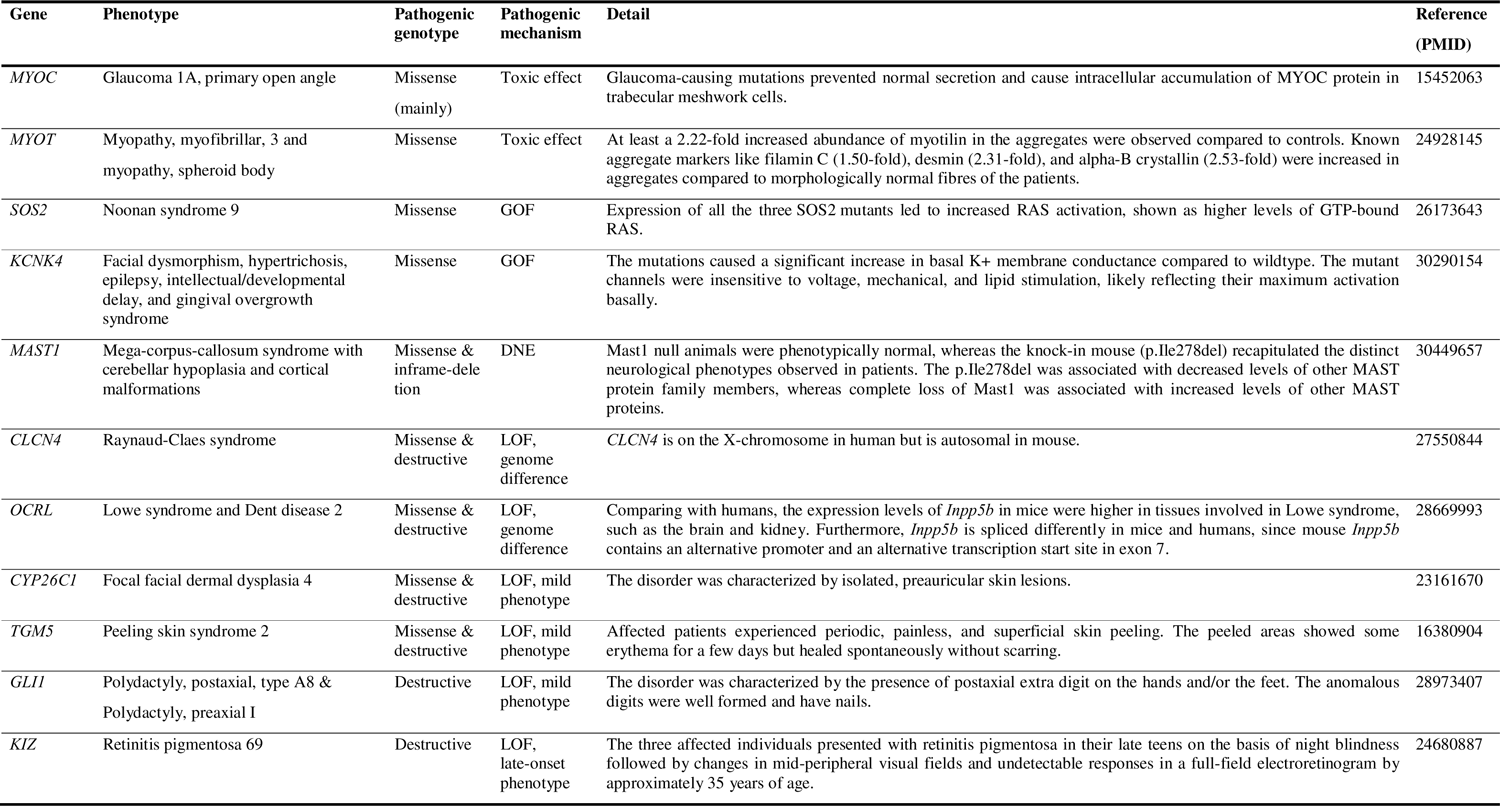
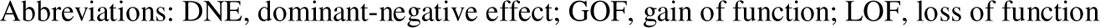
Pathogenic mechanisms of the dispensable/replaceable genes.

| Gene | Phenotype | Pathogenic genotype | Pathogenic mechanism | Detail | Reference (PMID) |
| --- | --- | --- | --- | --- | --- |
| *MYOC* | Glaucoma 1A, primary open angle | Missense (mainly) | Toxic effect | Glaucoma-causing mutations prevented normal secretion and cause intracellular accumulation of MYOC protein in trabecular meshwork cells. | 15452063 |
| *MYOT* | Myopathy, myofibrillar, 3 and myopathy, spheroid body | Missense | Toxic effect | At least a 2.22-fold increased abundance of myotilin in the aggregates were observed compared to controls. Known aggregate markers like filamin C (1.50-fold), desmin (2.31-fold), and alpha-B crystallin (2.53-fold) were increased in aggregates compared to morphologically normal fibres of the patients. | 24928145 |
| *SOS2* | Noonan syndrome 9 | Missense | GOF | Expression of all the three SOS2 mutants led to increased RAS activation, shown as higher levels of GTP-bound RAS. | 26173643 |
| *KCNK4* | Facial dysmorphism, hypertrichosis, epilepsy, intellectual/developmental delay, and gingival overgrowth syndrome | Missense | GOF | The mutations caused a significant increase in basal K<sup>+</sup> membrane conductance compared to wildtype. The mutant channels were insensitive to voltage, mechanical, and lipid stimulation, likely reflecting their maximum activation basally. | 30290154 |
| *MAST1* | Mega-corpus-callosum syndrome with cerebellar hypoplasia and cortical malformations | Missense & inframe-deletion | DNE | Mast1 null animals were phenotypically normal, whereas the knock-in mouse (p.Ile278del) recapitulated the distinct neurological phenotypes observed in patients. The p.Ile278del was associated with decreased levels of other MAST protein family members, whereas complete loss of Mast1 was associated with increased levels of other MAST proteins. | 30449657 |
| *CLCN4* | Raynaud-Claes syndrome | Missense & destructive | LOF, genome difference | *CLCN4* is on the X-chromosome in human but is autosomal in mouse. | 27550844 |
| *OCRL* | Lowe syndrome and Dent disease 2 | Missense & destructive | LOF, genome difference | Comparing with humans, the expression levels of *Inpp5b* in mice were higher in tissues involved in Lowe syndrome, such as the brain and kidney. Furthermore, *Inpp5b* is spliced differently in mice and humans, since mouse *Inpp5b* contains an alternative promoter and an alternative transcription start site in exon 7. | 28669993 |
| *CYP26C1* | Focal facial dermal dysplasia 4 | Missense & destructive | LOF, mild phenotype | The disorder was characterized by isolated, preauricular skin lesions. | 23161670 |
| *TGM5* | Peeling skin syndrome 2 | Missense & destructive | LOF, mild phenotype | Affected patients experienced periodic, painless, and superficial skin peeling. The peeled areas showed some erythema for a few days but healed spontaneously without scarring. | 16380904 |
| *GLI1* | Polydactyly, postaxial, type A8 & Polydactyly, preaxial I | Destructive | LOF, mild phenotype | The disorder was characterized by the presence of postaxial extra digit on the hands and/or the feet. The anomalous digits were well formed and have nails. | 28973407 |
| *KIZ* | Retinitis pigmentosa 69 | Destructive | LOF, late-onset phenotype | The three affected individuals presented with retinitis pigmentosa in their late teens on the basis of night blindness followed by changes in mid-peripheral visual fields and undetectable responses in a full-field electroretinogram by approximately 35 years of age. | 24680887 |

Abbreviations: DNE, dominant-negative effect; GOF, gain of function; LOF, loss of function

Taken together, the GDN is potentially associated with the pathogenic potential and pathogenic features of genes, including whether a gene is potentially pathogenic, the strength of pathogenicity that correlated with clinical phenotype spectrum (severe/mild phenotype or susceptibility), and the pathogenic genotype (mechanism) in some genes.

Previously, several parameters have been developed to predict the pathogenic potential of genes, including the McDonald-Kreitman neutrality index (NI) ^11^, residual variation intolerance score (RVIS) ^12^, gene damage index (GDI) ^13^, and LOF observed/expected upper bound fraction (LOEUF) ^14^. There were general correlations between the GDN and these parameters (Figure S1). However, the distributions of the scores of NI/RVIS/GDI/ LOEUF were obviously overlapped in genes of different GDNs. LOEUF presented more lineal correlation and less overlaps. However, using LOEUF <0.35 as a threshold of pathogenicity (suggested by gnomAD), only 32.9% vital genes were predicted to be intolerant to LOF variation. The LOEUF scores also varied much in the disease-associated genes, with only 23.8% of the genes associated with monogenic diseases having a LOEUF score <0.35 (Figure 1c, left panel). There was no significant correlation between LOEUF value and phenotype nature (severity). (Figure 1c, right panel).

### The GDQ and pathogenicity of variants/pathogenic feature of genes

From the perspective of genetic KO, the GDQs of 7,674 genes (72.7%), which showed abnormal phenotypes or embryonic/early death in the homozygous KO model but were asymptomatic in heterozygous KO, were estimated to be <50%; and the GDQs of 2,875 genes (27.3%) that showed abnormal phenotypes in heterozygous KO model were estimated to be >50% (Figure 2a).

**Figure 2.**
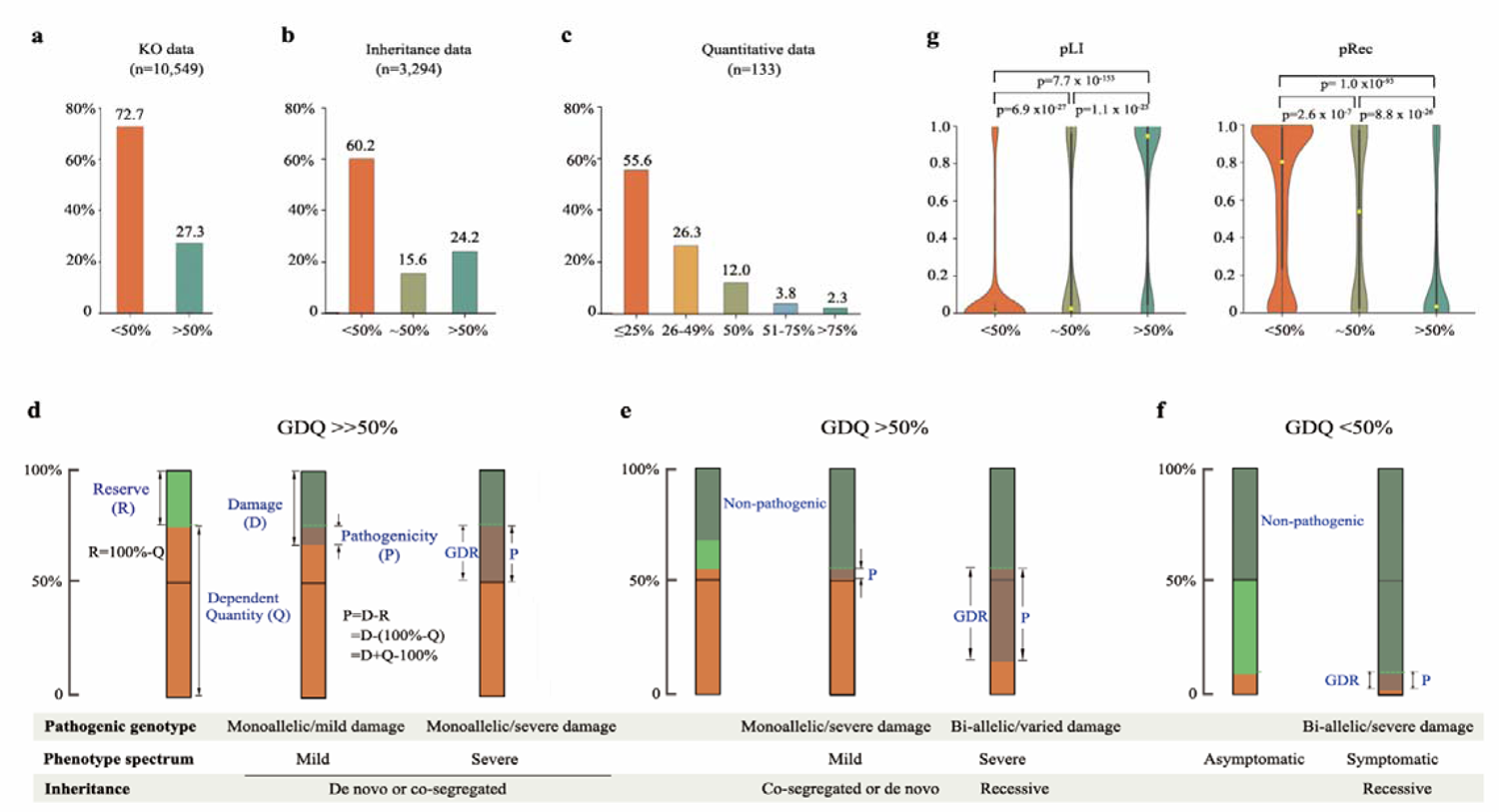
The genetic dependent quantity (GDQ) and the pathogenicity of genetic variant. (**a**) Distribution of GDQs determined by KO data. (**b**) Distribution of GDQs suggested by inheritance pattern. (**c**) Distribution of GDQs determined by quantitative experiments. (**d)** The model in a gene of high GDQ. The full scale of function of a gene in diploid creature was set as 100%. The essential quantity required for normal functioning (GDQ, Q) is less than 100%. The rest is considered as reserve (R=100%−Q). A variant is pathogenic when its damage (D) is large enough to run out of the reserve (R) and impaired the GDQ (Q, P=D-R=D+Q−100%). After exceeding the GDQ, the damages of variants are quantitatively correlated with the pathogenicity over a range (GDR) and potentially associated with a phenotype spectrum of varying severity. In a gene of GDQ >>50%, the monoallelic variants are pathogenic and associated a spectrum of phenotypes varying from mild to severe depending on the damage of variants. The variants would be originated de novo in individuals of severe phenotype, or co-segregated in a family with relative mild phenotype. (**e**) In genes of GDQ >50% slightly, both monoallelic and bi-allelic variants are potentially pathogenic. Monoallelic variants would commonly associate with mild cases; whereas bi-allelic variants would potentially be associated with severe phenotype. (**f)** In genes of GDQ <50%, which are common in diploids, only bi-allelic variants are pathogenic. The GDRs are relatively narrow in genes with very low GDQs. (**g**) Distributions and correlations between GDQ and the probability of being loss of function intolerant-pLI/pRec (left/right). The yellow circles mark the median, bold black bars indicate the 25^th^ to 75^th^ quartiles, and thick black bars indicate lower and upper boundaries of 1.5* or 25^th^ (75^th^) quartiles. Kruskal-Wallis test was used to estimate the differences among groups.

We then estimated the GDQs of disease-associated genes from the clinical-genetic perspective. We retrieved 4,268 genes that were associated with monogenic genetic disorders, among which 3,539 genes were classified as “pathogenic”/“possible pathogenic” in gene-disease association ^9^. After excluding genes with pathogenic mechanism other than LOF, a total of 3,294 genes were evaluated for their GDQs. The GDQs of 796 genes (24.2%), which were associated with diseases of dominant inheritance, were estimated to >50%; whereas the GDQs of 1,985 genes (60.2%), which were associated with diseases of recessive inheritance, were estimated to be <50%. Interestingly, 513 (15.6%) genes were associated with diseases of both dominant and recessive inheritances, and their GDQs were estimated to be ∼50% (Figure 2b).

Quantitative evidence from experiments allowed us to determine the precise GDQs for 133 genes, including 122 genes with residual activity data, 7 genes with mosaic evidence, and 5 genes with evidence from splicing analysis (note that *CFTR* had evidence for both residual activity and splicing analysis). The GDQs varied from 1.0% for *ADA* to 83% for *SCN1A*, including 8 genes with GDQ >50%, 16 genes with GDQ ∼50%, and 109 genes with GDQ <50% (Table S2). The distribution of the GDQs of these genes appeared as a continued spectrum with the genes of GDQ ≤25% being the most common (Figure 2c).

A variant is pathogenic when its damage exceeds the GDQ. After exceeding the GDQ, the damage of variants is quantitatively correlated with the pathogenicity within a range (genetic dependent range, GDR) (Figure 2d-f). Among the 133 genes with quantitative GDQ data available, the GDRs were defined for 75 genes (Table S2). Furthermore, 18 genes had well-defined GDQ and GDR (i.e., quantitative correlation in at least three levels and data on the upper and lower limit of the GDQ) (Table 2). The GDR of each gene also varied, from relatively wide to narrow (Table S2), which was consistent with the spectrum of phenotype that potentially ranged from susceptibility/mild phenotype (common) to severe genetic disease (rare).

**Table 2.**
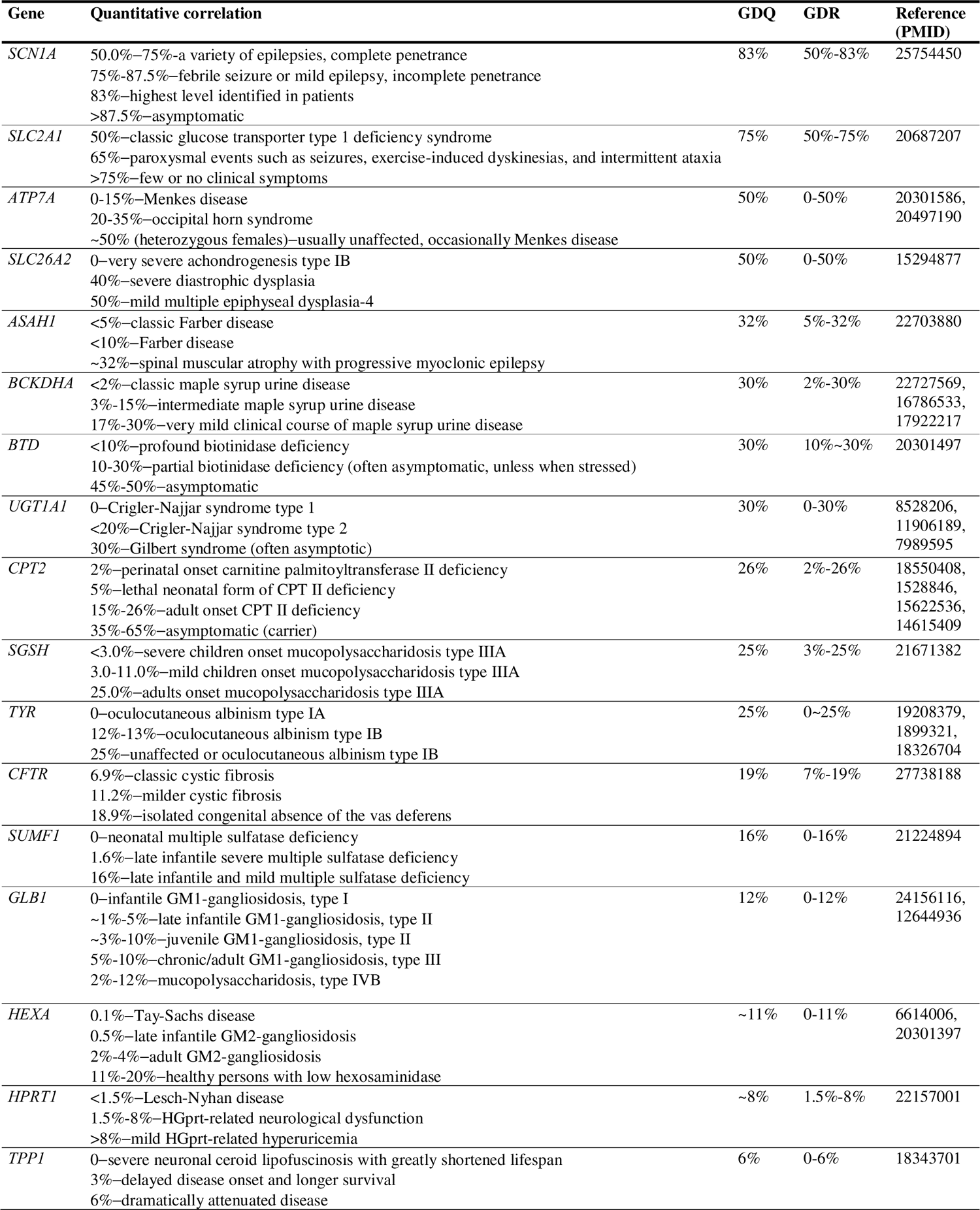
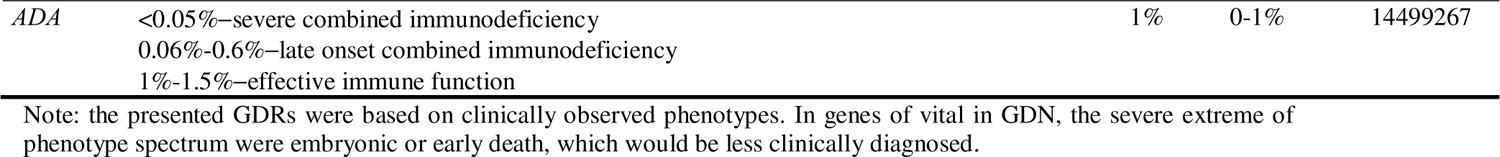
The genes with well defined genetic dependent quantity (GDQ) and genetic dependent range (GDR)

Since each gene differs in its GDQ, the pathogenicity (P) of the variants depends not only on the damage (D), but also on the GDQ (Q, P = D+Q–100%) (Figure 2d). The GDQ-damage relationship also determines the pathogenic features of the genes, including pathogenic genotype, phenotype spectrum, and inheritance of pathogenic variants (Figure 2d-f), which is the basis of evaluating the pathogenicity of variants in clinical practice ^15^.

Recently, the probability of being LOF intolerant (pLI/pRec) was proposed to predict whether genes are intolerant to heterozygous/homozygous variants ^14^. The genes with GDQs >50% were enriched significantly in the high pLI tail, whereas the genes with GDQs <50% were enriched significantly in the high pRec tail; and genes with GDQs ∼50% presented middle pLI and pRec values, although the distributions were bimodal (Figure 2g). These results suggest that the pLI and pRec, or similar parameters with compound heterozygous variants considered ^16^, could potentially be used to predict the GDQs.

### The GDS and clinical feature of genetic diseases

We analyzed gene expression levels across developmental stages and summarized the GDSs into five patterns (Figure 3a). Among the 17,846 genes with information of expressional stage available, 306 genes (1.7%) had a whole life stable pattern; and the genes of increasing pattern, decreasing pattern, “A” pattern, and “V” pattern accounted for 11.1%, 14.0%, 19.5%, and 53.7%, respectively (Figure 3b).

**Figure 3.**
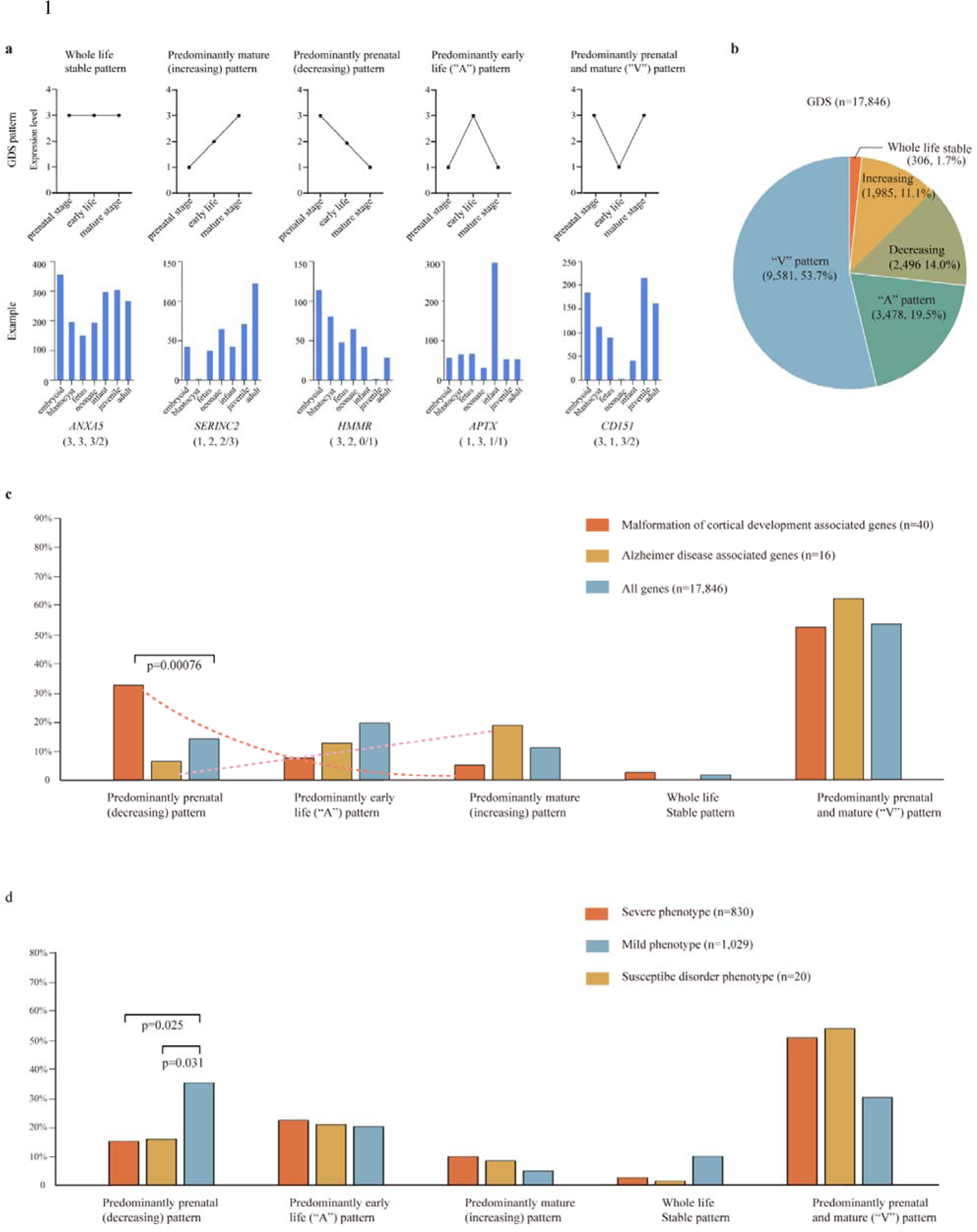
The genetic dependent stage (GDS) and the relationship between the GDS and the onset age/nature of genetic disorders. (a) **Classification of the GDS** pattern. A example of each pattern is illustrated in the bottom. **(b)** The distribution of GDS patterns in the 17,846 genes with expressional information available from the UniGene. (**c**) The correlation between the GDS and the genetic diseases of different onset age. Genes associated with malformation of cortical development had a significantly higher proportion of genes of GDS predominately in prenatal stages, and tended to have lower proportion of genes of GDS predominantly in mature stages. In contrast, the genes potentially associated with Alzheimer’s disease tended to have higher percentage of genes of GDS predominantly in mature stage. The two groups of disease-associated genes were obtained from the OMIM database. (**d**) The correlation between the GDS and the nature of genetic disorders (disease/susceptibility). In genes with validated pathogenic potential and GDN of vital, those associated with susceptibility had a significantly higher proportion of genes predominately expressed in prenatal stages (decreasing pattern), comparing with the genes associated with diseases (severe/mild). Chi-square statistical analysis was applied to compare the difference of frequencies in (**c**) and (**d**).

The GDS of genes is potentially associated with the onset age of genetic disorders. We analyzed genes associated with malformation of cortical development (including lissencephaly, microcephaly, and holoprosencephaly) and genes associated with Alzheimer’s disease (Figure 3c). The genes associated with malformation of cortical development had a significantly higher proportion of genes predominately in prenatal stages, while the genes potentially associated with Alzheimer’s disease tended to have higher percentage of genes predominantly in the mature stage.

The GDS is potentially associated with the phenotype spectrum (phenotype severity) and nature of genetic disorders (disease/susceptibility). For instance, *SCN1A* is expressed predominantly in the early life (1, 3, 0/1), which explain the early seizure-onset and the whole spectrum of phenotype with the severe extreme of Dravet syndrome. In contrast, *UNC13B* is also vital in GDN, but is expressed predominantly during prenatal and mature stage (3, 0, 1/2). *UNC13B* is therefore potentially associated with a phenotypical spectrum with the severe extreme of embryonic/early death that was less often identified clinically and a common mild epilepsy of late (even adult) onset ^17^ at the rest part of the spectrum of disease. Similarly, genes associated with susceptibility may be of GDS predominately in prenatal period, typically as *BRACA2* (3, 0, 0/1, GDN of vital), which is potentially associated with embryonic/early death in one extreme and susceptibility to cancer in the other part of phenotype spectrum ^18^. We therefore studied the correlation between GDS and nature of genetic diseases. Among the gens of GDN of vital, genes associated with susceptibility had a significantly higher proportion of genes predominately expressed in prenatal stages (decreasing pattern) (Figure 3d).

The GDS potentially determines the nature course of genetic disorders. Our recent study showed that seizure-free outcome in infant spasm patients with *ZFHX3* variants ^19^ and self-limited epilepsy in patients with *PCDH15* variants ^20^ were correlated with the GDS. The relapse of epilepsy after long-term of seizure-free in patients with *FAT1* variants was associated with the second expression peak of *FAT1* in adolescent and adult ^21^, which imply a significance in clinical practice like determining the optimal duration of therapy.

## Discussion

About 150 years ago, the experiments performed by Mr. Gregor Mendel formed the basis of genetics. Based on the genetics of Mendel, 3,539 genes have been identified to be associated with human diseases ^9^, mostly “rare diseases”. However, the classical Mendelian genetics has difficulties in explaining the inheritance of common phenotype/common diseases, typically shown by the facts that crossing of yellow-seeded plants with green-seeded ones produced either yellow or green peas, while more commonly offspring showed a mixed phenotype/trait of each parent. The mechanisms underlying the inheritance pattern and the pathogenicity of genes/variants remain unclear. The current major challenges in genetic studies and precision medicine include how to identify the causative genes of common diseases like genetic generalized epilepsy; how to determine the gene-disease associations, especially for some long-term refuted pathogenic genes such as *EFHC1* and *CACNA1H;* and how to evaluate the pathogenicity of variants in a given gene, as in the case of variants of *SLC2A1* (encoding GLUT1) ^3,7, 8^. This study shows each gene differs in genetic dependent features that determine the gene-disease associations extensively. The GDN determines whether a gene is potentially pathogenic, the strength of pathogenicity that correlated with clinical phenotype spectrum, and the pathogenic genotype (mechanism) in some genes. The GDQ determines the pathogenicity of variants and also the pathogenic features of genes that include inheritance pattern, pathogenic genotype, and phenotype spectrum. The GDS is associated with several clinical characteristics of genetic diseases, including the onset age, nature evolution course, and nature of genetic disorders (disease/susceptibility). The quantitative feature of genetic dependence, including the continuous distribution of GDQ of genome and the GDR, together with the varied GDN and GDS in each gene that correlated with the phenotype spectrum and nature of genetic disorders, discloses the genetic mechanisms underlying common diseases. These findings provided the theoretical bases for formulation of protocols to identify genetic causes of common diseases and guidelines to evaluate the pathogenicity of the genes and genetic variants.

This study showed that over 85% of the genes play vital or obligatory roles; while the fractional and dispensable/replaceable genes could also be associated with diseases via specific mechanisms or with specific phenotypes. However, the gene-disease associations in four fifth of human genome are not determined, despite the wide application of whole gene/exon sequencing on common diseases of large cohorts ^7^. To identify novel pathogenic genes associated with common diseases, several suggestions would be obtained by understanding the genetic dependent features.

First, the GDN and common diseases. The genes with GDN of obligatory are mostly common (44.5%). Comparing with the genes with GDN of vital, such genes are commonly associated with milder abnormal phenotype or susceptibility alteration, which would be more common than severe “rare” diseases due to the less pressure of natural selection on the phenotype-associated variants. Pathogenicity of the genes with GDN of obligatory tends to be moderate and should be evaluated from a comprehensive perspective of pathogenic feature ^9^, to avoid being in long-term dispute ^7^. Additionally, the genes with GDN of fractional/dispensable are also potentially associated with phenotypes that had less effect on human life, thus being mild and common.

Second, the GDN/GDS and common diseases. Genes with GDN of vital are also potentially associated with common milder phenotype or susceptibility, when the gene is expressed predominantly in prenatal stages (decreasing or “V” pattern), since severe impairment of such genes would result in embryonic/early death that is at the severe extreme of phenotype spectrum and would be less often identified clinically, and the mild phenotypes at the rest part of spectrum would be clinically common. We recently focused on common epilepsies with favorable outcomes. As a matter of fact, majority of the newly identified candidate epilepsy genes are of vital in GDN and of “V” pattern in GDS, including *AFF2* ^22^*, BCOR* ^23^*, CELSR1* ^24^*, CHD4* ^25^, *LAMA5* ^26^, *PKD1* ^27^, *UNC13B* ^17^, and *RYR2* ^28^.

Third, GDQ and inheritance feature/common diseases. De novo ^29^ or ultra-rare ^30^ variants have been targeted in recent studies to identify novel pathogenic genes. However, single pathogenic variants (de novo/ultra rare) appear only in genes with GDQ >50%, especially in those of high GDQ such as *SCN1A* (Table 2). In fact, genes of lower GDQ are common. The continuous distribution of GDQ suggests attention to variants of different inheritance pattern in designing protocols for identifying novel disease-associated genes, including de novo, bi-allelic, hemizygous, and co-segregated variants, which occur in genes of different GDQs. The novel epilepsy genes we identified recently in common epilepsies include genes of AD (*ATP6V0C* ^31^*, CELSR3* ^32^, *CHD4* ^25^, and *SBF1* ^16^), AR (*CELSR2* ^16^, *FAT1* ^21^, *LAMA5* ^26^, *MPDZ* ^33^, *PKD1* ^27^, and *ZFHX3* ^19^), X-linked recessive (*AFF2* ^22^*, BCOR* ^23^, *SHROOM4* ^34^, and *TENM1* ^16^), and AD & AR (*BSN* ^35^, *CELSR1* ^24^, and *UNC13B* ^17^). By employing trio-based individualized analysis, we recently identified three novel candidate genes associated with childhood epileptic encephalopathies. These genes included *SBF1* with de novo dominance pattern, *CELSR2* with recessive inheritance, and *TENM1* with X-linked recessive inheritance, which highlights the implication of trio-based strategies with emphases on different inheritance in identifying novel disease-causing genes of different GDQs ^16^.

Fourth, the GDQ/GDR and common diseases. Due to the quantity-dependent feature (GDQ and GDR), the phenotype caused by abnormalities in a given gene may vary in the severity that was correlated with the severity of damaging effect (Figure 2). Genetic variants with less damaging effect, such as missense, would be more common than variants with more severe damaging effect under the pressure of natural selection, subsequently leading to mild and common diseases, such as *SCN1A* variants that are associated with rare disease of Dravet syndrome but more commonly are associated with mild epilepsies like partial epilepsy with FS plus ^36^ or FS ^37^. Attention is required to variants with less damaging effect, besides the null variants ^38^, in genetic study as well as clinical practice in evaluating the pathogenicity of variants. In fact, most variants that we identified in common epilepsies are missense ^16, 17, 19, 21–27, 31–35^.

Fifth, the GDS and common diseases. This study demonstrated a potential correlation between the GDS and the onset age of genetic disorders. So far, development-related phenotypes ^9^, such as epileptic encephalopathies, are commonly studied (www.omim.org). However, many genes in human genome express in an increasing pattern (11.1%) or “V” pattern (53.7%), which are potentially associated with late-onset common genetic disorders, like *UNC13B* ^17^ (3, 0, 1/2, “V” pattern). The GDS is associated with evolution/outcome of genetic diseases, which is potentially one of the determinants of common diseases with favorable outcomes.

Regarding determining the pathogenic potential of genes, several parameters, such as NI, RVIS, GDI, and LOEUF, had been developed previously. These parameters are associated with the pathogenicity of genes in varied degrees. The present study shows that the genetic dependent feature determines the gene-disease association extensively, including whether a gene is pathogenic and the pathogenic features. With consideration on the genetic dependent feature, we developed a pathogenic potential and pathogenic feature assessment (PPA) framework to evaluate the gene-disease association with criteria on phenotype (spectrum) specificity, inheritance pattern, genotype-phenotype correlation, gene expressional profile, and gene KO consequence^9^. Besides the pathogenic potential, the pathogenic features of each gene were evaluated (www.gdap.org.cn), which are essential part of the gene-disease association and critically useful in evaluating the pathogenicity of variants of the genes ^15^.

On evaluating the pathogenicity of variants, the damaging effects of variants are commonly considered previously ^38^, but the GDQs of the genes are usually not taken into account. Theoretically, the pathogenicity (P) of a variant could be determined by quantifying the damage (D) of the variant and the GDQ (Q) of the gene (P=D+Q–100%, Figure 2). However, quantifying the GDQ of each gene and the damage of each variant appears impracticable. Based on the GDQ-damage relationship that determines the pathogenic genotype, phenotype spectrum, and inheritance of pathogenic variants, we developed a clinical concordance evaluation (CCE) framework to evaluate the pathogenicity/causality of variants ^15^. The CCE is an individualized protocol to compare the clinical feature of a patient with the pathogenic feature of a candidate gene from four aspects, which considered not only the damaging effects of variants as previous protocols ^38^, but also the GDQ that differs in each gene. Potential advantages of CCE in specificity and sensitivity were suggested by its clinical application ^15^. The results of PPA, together with the data of genetic dependence and other information required for CCE (www.gdap.org.cn), provided the bases for individualized clinical diagnosis of genetic diseases.

This study suggests that each gene differs in genetic-dependent feature that determines the gene-disease association extensively. However, future studies are required to elaborate the genetic dependent features. First, nearly half of the genes in the human genome lack information of the GDN; and the current evidence on GDN were primarily from genetic KO animals, and mostly mice, which should be interpreted with understanding the difference between human and animal and among different animal strains. Second, the GDQs of most genes remain to be specified. Third, further studies are required to determine the precise temporal patterns of gene expression, including the tissue-specific expression pattern as indicated in our recent study on *ZFHX3* ^19^ and *PCDH15* ^20^.

In conclusion, the genetic dependence, potentially as a principle of genetics, discloses the genetic mechanisms of common diseases and the pathogenicity of genes/variants. The genetic dependent features of genes determine whether a gene is pathogenic, the strength of pathogenicity, and pathogenic features of the gene and several critical clinical features of the genetic disorder, such as the inheritance pattern, pathogenic genotype, phenotype spectrum, and onset/evolution/outcome of the genetic disorder. Understanding how human life depends on the genes is essential to know the gene-disease association. The increased characterizing of genetic dependent feature of genome, together with the novel discovery and determination of gene-disease associations ^9^ and the clinical application of individualized evaluation of sequence variants ^15^, would establish the fundamental bases of precise medicine of the future.

### Data access

Not applicable

### Data Availability

All data are available in the main text, the supplementary materials, and the GD&P database (www.gdap.org.cn).

## Acknowledgments

We thank Wen-Jun Zhang, Li-Hong Liu. Xiao-Yu Liang, Mi Jian, De-Hai Liang, Juan Wang, Zi-Long Ye, Nan-Xiang Shen, Xue-Qing Tang, and Huan Li for the pathogenic potential and pathogenic feature assessment. We thank Liang-Di Gao, Jiang-Guo Zhang, Yu-Lan Chen, Juldiz Ershn, and Qing Zhou for contribution in establishing the GD&P database.

## Author’s contributions

LWP conceived and designed the study, analyzed the data and wrote the paper. LB, BWJ and ZP collected and analyzed the data, performed statistical analysis. WJ, FCX XHQ and YL collected and analyzed the data. HN, SYW, ST, and YYH performed data analysis and interpretation.

## Additional Information

Figure S1-S2 Table S1-S2

## References

1. Ezkurdia I, Juan D, Rodriguez JM, et al. Multiple evidence strands suggest that there may be as few as 19,000 human protein-coding genes. Human molecular genetics 2014; 23(22): 5866–78.

2. Couzin J. To what extent are genetic variation and personal health linked? Science 2005; 309(5731): 81.

3. MacArthur DG, Manolio TA, Dimmock DP, et al. Guidelines for investigating causality of sequence variants in human disease. Nature 2014; 508(7497): 469–76.

4. Breschi A, Gingeras TR, Guigo R. Comparative transcriptomics in human and mouse. Nature reviews Genetics 2017; 18(7): 425–40.

5. Shi YW, Yu MJ, Long YS, et al. Mosaic SCN1A mutations in familial partial epilepsy with antecedent febrile seizures. Genes, brain, and behavior 2012; 11(2): 170–6.

6. Clavero S, Perez B, Rincon A, Ugarte M, Desviat LR. Qualitative and quantitative analysis of the effect of splicing mutations in propionic acidemia underlying non-severe phenotypes. Human genetics 2004; 115(3): 239–47.

7. McKee JL, Karlin A, deCampo D, Helbig I. GLUT1, GGE, and the resilient fallacy of refuted epilepsy genes. Seizure 2023; 109: 97–8.

8. He N, Li B, Lin ZJ, Zhou P, Su T, Liao WP. Common genetic epilepsies, pathogenicity of genes/variants, and genetic dependence. Seizure 2023; 109: 38–9.

9. Bian WJ, Wang J, Li B, et al. Gene-disease association: pathogenic potential/pathogenic feature assessment.

10. Depienne C, Bouteiller D, Keren B, et al. Sporadic infantile epileptic encephalopathy caused by mutations in PCDH19 resembles Dravet syndrome but mainly affects females. PLoS genetics 2009; 5(2): e1000381.

11. McDonald JH, Kreitman M. Adaptive protein evolution at the Adh locus in Drosophila. Nature; 351(6328): 652–4.

12. Petrovski S, Wang Q, Heinzen EL, Allen AS, Goldstein DB. Genic intolerance to functional variation and the interpretation of personal genomes. PLoS genetics 2013; 9(8): e1003709.

13. Itan Y, Shang L, Boisson B, et al. The human gene damage index as a gene-level approach to prioritizing exome variants. Proceedings of the National Academy of Sciences of the United States of America 2015; 112(44): 13615–20.

14. Karczewski KJ, Francioli LC, Tiao G, et al. The mutational constraint spectrum quantified from variation in 141,456 humans. Nature 2020; 581(7809): 434–43.

15. Zhou P, He N, Lin Z-J, et al. Clinical concordance evaluation of the causality of sequence variants.

16. Shi Y-W, Zhang J, He N, et al. Variants in SBF1, CELSR2, and TENM1 are associated with childhood epileptic encephalopathies. *medRxiv* 2023: 2023.07.25.23293037.

17. Wang J, Qiao JD, Liu XR, et al. UNC13B variants associated with partial epilepsy with favourable outcome. Brain: a journal of neurology 2021; 144(10): 3050–60.

18. Narod SA, Foulkes WD. BRCA1 and BRCA2: 1994 and beyond. Nat Rev Cancer 2004; 4(9): 665–76.

19. He M-F, Liu L-H, Luo S, et al. ZFHX3 Associated with Partial Epilepsy/Spasms and Correlation between Outcome & Gene Expression Stage. medRxiv 2023: 2023.07.16.23292551.

20. Lin S-M, Su T, Qiao J-D, et al. PCDH15 in self-limited partial epilepsy and the mechanism of distinct phenotypes. 2023.

21. Zou DF, Li XY, Lu XG, et al. Association of FAT1 with focal epilepsy and correlation between seizure relapse and gene expression stage. Seizure 2023.

22. Zou D, Qin B, Wang J, et al. AFF2 Is Associated With X-Linked Partial (Focal) Epilepsy With Antecedent Febrile Seizures. Frontiers in Molecular Neuroscience 2022; 15.

23. Li X, Bian W-J, Liu X-R, et al. BCOR variants are associated with X-linked recessive partial epilepsy. Epilepsy Research 2022; 187: 107036.

24. Chen Z, Luo S, Liu ZG, et al. CELSR1 variants are associated with partial epilepsy of childhood. American Journal of Medical Genetics Part B: Neuropsychiatric Genetics 2022; 189(7-8): 247–56.

25. Liu XR, Ye TT, Zhang WJ, et al. CHD4 variants are associated with childhood idiopathic epilepsy with sinus arrhythmia. CNS neuroscience & therapeutics 2021; 27(10): 1146–56.

26. Luo S, Liu Z-G, Wang J, et al. Recessive LAMA5 Variants Associated With Partial Epilepsy and Spasms in Infancy. Frontiers in Molecular Neuroscience 2022; 15.

27. Wang J-Y, Wang J, Lu X-G, et al. Recessive PKD1 Mutations Are Associated With Febrile Seizures and Epilepsy With Antecedent Febrile Seizures and the Genotype-Phenotype Correlation. Frontiers in Molecular Neuroscience 2022; 15.

28. Ma MG, Liu XR, Wu Y, et al. RYR2 Mutations Are Associated With Benign Epilepsy of Childhood With Centrotemporal Spikes With or Without Arrhythmia. Frontiers in neuroscience 2021; 15: 629610.

29. He N, Lin ZJ, Wang J, et al. Evaluating the pathogenic potential of genes with de novo variants in epileptic encephalopathies. Genetics in medicine: official journal of the American College of Medical Genetics 2019; 21(1): 17–27.

30. Chen S, Neale BM, Berkovic SF. Shared and distinct ultra-rare genetic risk for diverse epilepsies: A whole-exome sequencing study of 54,423 individuals across multiple genetic ancestries. medRxiv 2023.

31. Tian Y, Zhai QX, Li XJ, et al. ATP6V0C Is Associated With Febrile Seizures and Epilepsy With Febrile Seizures Plus. Front Mol Neurosci 2022; 15: 889534.

32. Li J, Lin SM, Qiao JD, et al. CELSR3 variants are associated with febrile seizures and epilepsy with antecedent febrile seizures. CNS neuroscience & therapeutics 2022; 28(3): 382–9.

33. Luo J, Li Y, Lv Y, et al. MPDZ variants associated with epilepsies and/or febrile seizures and the individualized genotype-phenotype correlation. Genes & Diseases 2023.

34. Bian WJ, Li ZJ, Wang J, et al. SHROOM4 Variants Are Associated With X-Linked Epilepsy With Features of Generalized Seizures or Generalized Discharges. Front Mol Neurosci 2022; 15: 862480.

35. Ye T, Zhang J, Wang J, et al. Variants in BSN gene associated with epilepsy with favourable outcome. Journal of Medical Genetics 2022: jmg-2022–108865.

36. Liao WP, Shi YW, Long YS, et al. Partial epilepsy with antecedent febrile seizures and seizure aggravation by antiepileptic drugs: associated with loss of function of Na(v) 1.1. Epilepsia 2010; 51(9): 1669–78.

37. Skotte L, Fadista J, Bybjerg-Grauholm J, et al. Genome-wide association study of febrile seizures implicates fever response and neuronal excitability genes. Brain: a journal of neurology 2022; 145(2): 555–68.

38. Richards S, Aziz N, Bale S, et al. Standards and guidelines for the interpretation of sequence variants: a joint consensus recommendation of the American College of Medical Genetics and Genomics and the Association for Molecular Pathology. Genetics in medicine: official journal of the American College of Medical Genetics 2015; 17(5): 405–24.

