## Supplementary material for "Genetic Dependence and Genetic Diseases": Supplmentary figures and tables

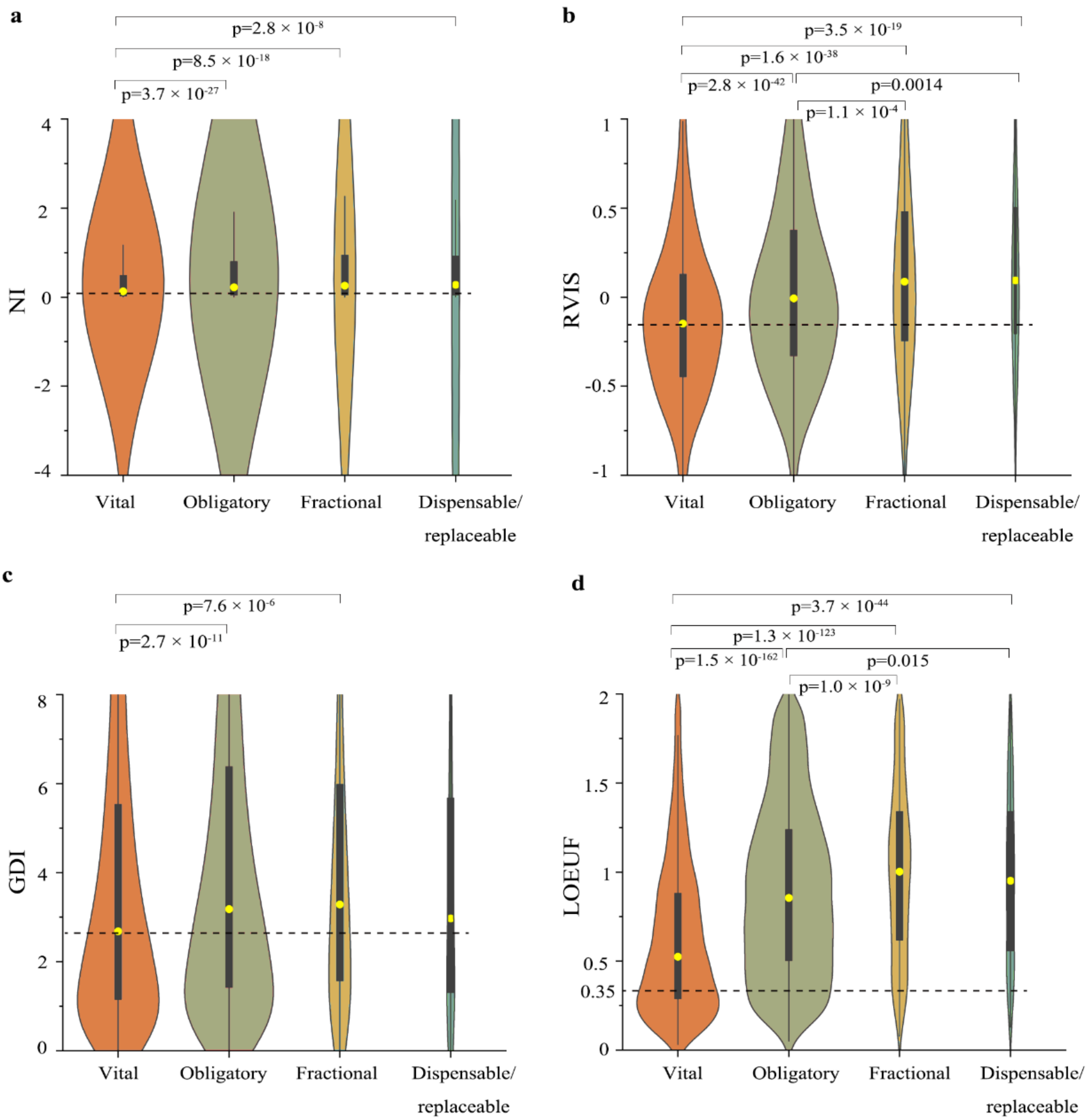


**Figure S1. Correlations between genetic dependent nature (GDN) and the parameters related to the pathogenic potential of genes. (a)** GDN and neutrality index (NI). Vital genes presented significantly lower median NI score than the other three groups. **(b)** GDN and residual variation intolerance score (RVIS). Vital genes presented significantly lower median RVIS score than the other three groups; the median RVIS score of obligatory genes was also significantly lower than that of fractional and dispensable/replaceable genes. **(c)** GDN and gene damage index (GDI). Vital genes presented significantly lower median GDI score than the obligatory and fractional genes. **(d)** GDN and loss of function (LOF) observed/expected upper bound fraction (LOEUF). Vital genes presented significantly lower median LOEUF score than the other three groups; the median LOEUF score of obligatory genes was also significantly lower than that of fractional and dispensable/replaceable genes. Gens with LOEUF <0.35 were predicted to be highly intolerant to LOF variations. The yellow circles marks the median, bold black bars indicate the $25^{th}$ to $75^{th}$ quartiles, and thick black bars indicate lower and upper boundaries of 1.5* or $25^{th}$ ($75^{th}$) quartiles. Kruskal-Wallis test was used to estimate the differences among groups.

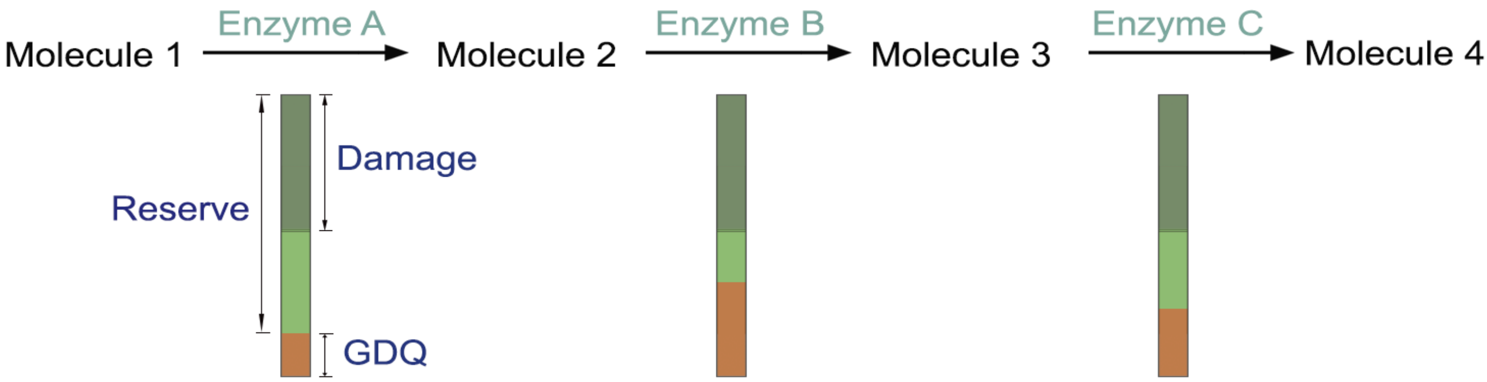


**Figure S2. The genetic dependent quantity (GDQ) explaining whether damages in different genes are addable.** Due to the feature of GDQ, variants that did not damage the GDQ would affect reserve only. In a metabolism pathway, for instance, if the gene encoding enzyme A, B, and C, respectively, had damages that did not impair the GDQ individually, each step will be passed successfully. No effect in concert will be presented. Using next generation sequencing technologies, thousands of variants could been detected in an individual, who maintains normal due to the protection offered by the mechanism of dependent quantity/reserve.

**Table S1. Pathogenic mechanisms of the fractional genes (n=64)**

| Gene | Knock-out phenotype | Phenotype | Pathogenic genotype | Pathogenic mechanism | Detail | Reference (PMID) |
|---|---|---|---|---|---|---|
| **1. Pathogenic mechanism other than LOF (n=32)** | | | | | | |
| *CADM3* | Delayed myelination | Charcot-Marie-Tooth disease type 2FF | Missense | Toxic effect | In Schwann cells, the mutant protein was retained in the ER and triggered the unfolded protein response. | 33889941 |
| *CEL* | Abnormal intestinal lipid absorption | Maturity-onset diabetes of the young, type VIII | Deletions in the GC-rich variable number of tandem repeats | Toxic effect | A high tendency of both intracellular and extracellular aggregate formation were observed, which was not seen for the normal protein and which led to the stimulation of the unfolded protein response. The altered protein region also contained a high number of cysteine residues with a potential for disulfide bridge formation. | 29233499 |
| *CHCHD10* | Mild mitochondrial respiration anomalies in skeletal muscle | *CHCHD10*-related disorders | Missense (mainly) | Toxic effect | Accumulation of TAR DNA-binding protein 43 in cell bodies of neurons in affected regions was often observed, indicating that a unifying pathology underlined the disease. Mitochondrial mis-localization of TAR DNA-binding protein 43 and its involvement in neurotoxicity of amyotrophic lateral sclerosis and frontotemporal lobe dementia had also been reported. | 30791515 |
| *COL8A2* | Thinner Descemet's membrane of the cornea | Corneal dystrophy, Fuchs endothelial, 1 | Missense | Toxic effect | Ultrastructural analysis showed the predominant effect to be dilated ER, suggesting ER stress and unfolded protein response (UPR) activation. Immunohistochemistry, Western blot, QT-PCR, and TUNEL analyses supported UPR activation and UPR-associated apoptosis in the mutant corneal endothelium. | 22002996 |
| *COMP* | Abnormal vascular wound healing | Epiphyseal dysplasia, multiple, 1 | Missense & inframe-insertion/deletion | Toxic effect | Mutations resulted in the misfolding of the mutated protein and its retention in the rough endoplasmic reticulum (rER) of chondrocytes. This protein retention resulted in ER stress that ultimately caused increased cell death in vitro. | 20301660 |
| *LYZ* | Lacked a detectable leukocytic isozyme | Amyloidosis, renal | Missense | Toxic effect | Mutations responsible for amyloid formation in immunoglobulin light chain and transthyretin variants affected the stability of the proteins and their tendency to aggregate. | 17269695 |

| | | | | | | |
|---|---|---|---|---|---|---|
| *MUC1* | Abnormal gallbladder physiology, abnormal intestinal lipid absorption, decreased intestinal cholesterol absorption, and gallstones. | Medullary cystic kidney disease 1 | A cytosine insertion in an extremely long GC-rich tandem repeat | Toxic effect | Co-staining of patient and control tissue additionally with antibodies against normal MUC1 demonstrated the specificity of the MUC1-fs (the name for the predicted mutant protein) antibodies for the mutant protein, with diffuse and/or fine granular intracellular localization of the MUC1-fs protein in patient kidney. | 23396133 |
| *NIPA1* | Decreased circulating high-density lipoprotein cholesterol level | Spastic paraplegia 6, autosomal dominant | Missense | Toxic effect | Mutant NIPA1 accumulated in the ER triggering ER stress and features of apoptotic cell death. | 19091982 |
| *TGM6* | Decreased circulating cholesterol level | Spinocerebellar ataxia 35 | Missense (mainly) | Toxic effect | Wild-type protein mainly localized to the nucleus and perinuclear area, whereas mutations showed nuclear depletion, increased accumulation in the perinuclear area, insolubility and loss of enzymatic function. | 28934387 |
| *FGF12* | Decreased thigmotaxis and abnormal coat appearance | Epileptic ence-phalopathy, early infantile, 47 | Missense | GOF | Functional studies showed enhanced modulation of channel inactivation gating, equivalent to a GOF effect. | 27164707 |
| *HRAS* | Chylous ascites and decreased incidence of tumors by chemical induction | Costello syndrome | Missense & inframe-insertion | GOF | The amino acid changes led either to decreased GTPase activity (if amino acids 12, 13, 59, 61, 63 were involved) so that oncogenic RAS mutant proteins were locked in the active GTP-bound state, or decreased nucleotide affinity, and hence, demonstrated increased exchange of bound GDP for cytosolic GTP. | 20301680 |
| *KCNT1* | Impaired action potential firing in sensory neurons and increased mechanical hypersensitivity in neuropathic pain models | Epilepsy, nocturnal frontal lobe, 5 | Missense & inframe-deletion | GOF | Functional consequences of mutations displayed a strong GOF effect by producing large-fold increases in the current amplitude. | 23086397 |
| *MRAS* | Decreased mean percentage of peripheral blood B cells | Noonan syndrome 11 | Missense | GOF | Biochemical studies showed a 40-fold increase in GTP loading, as well as increased extracellular signal-regulated kinase activation and signaling and transcription activation in response to growth factors. | 28289718 |
| *PIK3R2* | Lower blood glucose levels | Megalencephaly-polymicrogyria-polydactyly-hydrocephalus syndrome 1 | Missense | GOF | The recurrent pathogenic variant, p.G373R, caused hyperactivation of PI3K-AKT-MTOR pathway downstream targets. | 22729224 |

| | | | | | | |
|---|---|---|---|---|---|---|
| *PPP2R2B* | Abnormal circulating potassium level, decreased mean corpuscular hemoglobin, decreased mean corpuscular volume, and thrombocytopenia | Spinocerebellar ataxia 12 | CAG trinucleotide repeat expansion | GOF | Expanded CAG repeats in the 5-prime region of the *PPP2R2B* gene caused increased gene expression. | 20533062 |
| *PSTPIP1* | Defects in immune cells with T cell hyperresponsive to antigen receptor stimulation | Pyogenic sterile arthritis, pyoderma gangrenosum, and acne | Missense (mainly) | GOF | Mutations all fell within the coiled-coil region of the molecule and disrupted PSTPIP-1-PEST phosphatase binding and regulation, resulting in hyperphosphorylation of the mutant protein and increased avidity for pyrin. | 22161697 |
| *PTDSS1* | Decreased phosphatidylethanol-amine and phosphatidylserine levels in the liver, and increased circulating calcium level and prepulse inhibition | Lenz-Majewski hyperostotic dwarfism | Missense | GOF | Phosphatidylserine synthesis was increased in intact fibroblasts from affected individuals, and end-product inhibition of PSS1 by phosphatidylserine was markedly reduced. Therefore, mutations caused a GOF effect associated with regulatory dysfunction of PSS1. | 24241535 |
| *RIT1* | Increased cellular sensitivity to hydrogen peroxide | Noonan syndrome 8 | Missense | GOF | Compared with the WT cDNA, five *RIT1* mutations (p.S35T, p.A57G, p.Glu81Gly, p.F82L, and p.G95A) exhibited significant activation. | 23791108 |
| *RNF13* | Abnormal sleep behavior, abnormal hindlimb morphology, and abnormal limb position | Epileptic ence-phalopathy, early infantile, 73 | Missense | GOF | Both IRE1-alpha-mediated stress signaling and stress-induced apoptosis were increased in affected individuals' cells, indicating that the *RNF13* variants conferred GOF to the encoded protein and thereby led to altered signaling of the ER stress response associated with severe neurodegeneration in infancy. | 30595371 |
| *SH3BP2* | Higher pre-B cell numbers and impaired B cell receptor signaling or thymus-independent type 2 humoral responses | Cherubism | Missense (mainly) | GOF | Mutant myeloid cells showed increased responses to M-CSF and RANKL stimulation, and formed macrophages that expressed high levels of TNF-alpha and osteoclasts that were unusually large. | 11381256 |
| *SLC25A24* | Increased grip strength | Fontaine progeroid syndrome | Missense | GOF | Individuals with Fontaine syndrome showed normal amounts of SLC25A24 in fibroblasts but marked changes in mitochondrial morphology and function, suggesting a pathological GOF of the mutated SLC25A24. | 29100094 |
| *SNTA1* | Impaired astrocyte and neuromuscular synapse morphology | Long QT syndrome 12 | Missense | GOF | Electrophysiologic analysis suggested that A257G mutant channels exhibited a GOF through 3 mechanisms: increase of channel availability by leftward shift of activation kinetics, delay of current decay, and increase in current density. | 19684871 |

| | | | | | | |
|---|---|---|---|---|---|---|
| *TRPM4* | Abnormal dendritic cell physiology, increased vascular permeability, and increased susceptibility to type I hypersensitivity reaction | Cardiac conduction disorders | Missense | GOF | Mutation attenuated deSUMOylation of the TRPM4 channel, resulting in impaired endocytosis and elevated TRPM4 channel density at the cell surface. Electrophysiologic analysis indicated that mutations represented GOF. | 30528822 |
| *CRYBB3* | Abnormal epididymis morphology, enlarged epididymis, and increased bone mineral content | Cataract 22 | Missense & lengthened mutation | DNE | The mutation was predicted to cause destabilization of the domain structure of the Greek key motif IV and to affect CRYBB3 protein folding. | 23508780 |
| *GLDN* | Abnormal axon morphology | Lethal congenital contracture syndrome 11 | Missense & destructive | DNE | Pathogenic variants detected affected gliomedin's transportation to the cell surface and it's binding to NF186. | 35806855 |
| *HSPB1* | Impaired wound healing with delayed wound closing and abnormal inflammatory response | *HSPB1*-related disorders | Missense (mainly) | DNE | Stable truncated HSPB1 bound wild type HSPB1, and strongly impaired the ability of the cells to cope with unfolded protein stress, suggesting a DNE. | 26675522 |
| *KCNQ3* | Abnormal apamin-insensitive afterhyperpolarization currents in granule cells | Seizures, benign neonatal, type 2 | Missense (mainly) | DNE | Most variants caused a 20%-40% reduction in the maximal current of heteromeric Q2/Q3 channels, but expression of p.Y309R reduced heteromeric channel function by about 60%, suggesting a DNE. | 25524373 |
| *KRT6A* | Delayed wound healing | Pachyonychia, congenital 3 | Missense (mainly) | DNE | The mutant polypeptide was very different in both sequence and predicted secondary structure from the wild-type K6a tail domain, consistent with a DNE mutation. | 21326300 |
| *PHACTR1* | Decreased circulating triglyceride level and increased bone mineral content | Epileptic ence-phalopathy, early infantile, 70 | Missense | DNE | The mutant P.R521C competitively inhibited the function of reduced levels of endogenous PHACTR1, and thus exerted DNE. | 30256902 |
| *SGMS2* | Increased ceramide levels, decreased sphingomyelin, sphingomyelin-1-phosphate, and diacylglycerol levels, and resistance to lysenin-mediated cytolysis | Calvarial doughnut lesions with bone fragility with or without spondylometaphyseal dysplasia | Missense & frameshift | DNE | Robust expression of a 49-residue-long N-terminal SMS2 fragment may have a DNE on the bone regulatory properties of the full-length protein. | 30779713 |
| *TRIM8* | Increased NK cell number, abnormal homeostasis, and decreased bone mineral density | Focal segmental glomerulosclerosis and neurodevelopmental syndrome | Truncation | DNE | Variants may cause DNE of the wild-type allele, given that TRIM8 can dimerize through its coiled-coil domain27 and C-terminal case-associated variants cause TRIM8 mislocalization. | 33508234 |

| | | | | | | |
|---|---|---|---|---|---|---|
| *PCDH19* | Abnormal electrocorticograms and distinct clusters of null and wild-type cells in the brain | Epileptic ence-phalopathy, early infantile, 9 | Missense & destructive | Cellular interference | Heterozygous females (common) and mosacism males (rare) were pathogenic. Two populations of cells, mutant and wildtype, were required for disease to occur, whereas a homogeneous cell population, either mutant or wildtype, did not lead to disease. | 19214208 |
| **2. Pathogenic mechanism of LOF but associated with mild phenotype (n=18)** | | | | | | |
| *ACADSB* | Increased lean body mass | 2-methyl-butyrylglycinuria | Missense & destructive | LOF | The disorder was characterized by impaired isoleucine degradation. It was most often ascertained via newborn screening and was usually clinically asymptomatic. | 17945527 |
| *CAT* | Subtle abnormalities in mitochondrial respiration | Acatalasemia | Missense &destructive | LOF | The disorder was usually asymptomatic, sometimes oral ulcerations and gangrene may be present. | 24522161 |
| *CDH3* | Alveolar hyperplasia, ductal dysplasia, and extensive lymphocyte infiltration of the mammary glands | *CDH3*-related disorders | Missense &destructive | LOF | *CDH3*-related disorders were skin diseases characterized by hypotrichosis and macular dystrophy with variable additional limb and ectodermal anomalies. | 27386845 |
| *DNASE1L3* | Abnormal cell death | Systemic lupus erythematosus 16 | Destructive | LOF | Variants in this gene were associated with susceptibility to autoimmunity, such as systemic lupus erythematosus. The patho-aetiology probably depended on complex multifactorial interactions between various genetic, enzymatic, and environmental factors as well as on epistatic effects. | 18486922 |
| *EXPH5* | Abnormal circulating ion level | Epidermolysis bullosa, nonspecific, autosomal recessive | Destructive | LOF | The disorder was characterized by blistering of skin and mucosae, following minimal pressure or trauma. | 28830826 |
| *FGF5* | Longer pelage hair | Trichomegaly | Missense & destructive | LOF | Trichomegaly was characterized by unusually long eyelashes and mild hypertrichosis of eyebrow. | 24989505 |
| *FGF17* | Tissue losses in the inferior colliculus and the anterior vermis of the brain | Hypogonadotropic hypogonadism 20 with or without anosmia | Missense | LOF | The disorder was characterized by abnormal pubertal development and infertility. | 23643382 |
| *FZD6* | Abnormal hair follicle orientation | Nail disorder, nonsyndromic congenital, 1 | Missense & destructive | LOF | The disorder was characterized by excessive longitudinal striations and numerous superficial pits on the nails, which had a distinctive rough sandpaper-like appearance. | 23374899 |

| | | | | | | |
|---|---|---|---|---|---|---|
| *GALK1* | Tissue accumulation of galactose and galactitol but do not form cataracts | Galactokinase deficiency with cataracts | Missense &destructive | LOF | The disorder was characterized by early onset of cataracts but otherwise experience few long-term complications. | 22632133 |
| *ISG15* | Decreased bone mineralization, ossification and strength | Immunodeficiency 38 | Destructive | LOF | Individuals predisposed to severe clinical disease upon infection with weakly virulent mycobacteria but do not experienced severe disease in response to viral infection. | 22859821 |
| *LPAR6* | Increased mean corpuscular hemoglobin | *LPAR6*-related disorders | Missense & destructive | LOF | *LPAR6*-related disorders were rare genetic hair loss disorders characterized by sparse scalp hair, sparse to absent eyebrows and eyelashes, and sparse axillary and body hair. | 25828854 |
| *PLA2G5* | Reduced acute zymosan-induced peritonitis and arachadonic acid metabolite release from stimulated peritoneal macrophages | Fleck retina, familial benign | Missense &destructive | LOF | The disorder was characterized by distinctive retinal appearance and no apparent visual or electrophysiological deficits. Affected individuals were asymptomatic. | 22137173 |
| *SLC39A5* | Altered zinc homeostasis and increased susceptibility to zinc-induced pancretitis | Myopia 24, autosomal dominant | Missense & destructive | LOF | Myopia was a refractive error of the eye. Light rays from a distant object were focused in front of the retina and those from a near object were focused in the retina, therefore distant objects were blurry and near objects were clear. | 24891338 |
| *SULT2B1* | Abnormal cholesterol level | Ichthyosis, congenital, autosomal recessive 14 | Missense & destructive | LOF | The disorder was characterized by abnormal skin scaling over the whole body. | 28575648 |
| *TBXAS1* | Increased bleeding time and decreased platelet response to arachidonic acid | Ghosal heamtodiaphyseal dysplasia | Missense | LOF | The disorder was characterized by abnormal thick bones and a shortage of red blood cells. | 18264100 |
| *TMC6* | Decreased body weight and decreased circulating insulin level | Epidermodysplasia verruciformis | Destructive | LOF | The disorder was a rare genodermatosis associated with a high risk of skin cancer. It resulted from an abnormal susceptibility to specific related human papillomavirus. | 10844558 |
| *TMEM126A* | Decreased circulating low density lipoprotein cholesterol level and decreased T-helper cell number | Optic atrophy-7 | Missense & destructive | LOF | Optic atrophy was characterized by bilateral deficiency in visual acuity, optic disk pallor, and central scotomas. | 30961538 |
| *ZNF341* | Decreased circulating insulin level, increased circulating chloride level, and increased circulating sodium level | Hyper-IgE recurrent infection syndrome 3, autosomal recessive | Destructive | LOF | The disorder was characterized by childhood onset of atopic dermatitis, skin infections particularly with Staphylococcus aureus, recurrent sinopulmonary infections, and increased serum IgE and IgG. | 29907691 |

**3. Pathogenic mechanism of LOF but associated with late-onset phenotype (n=9)**

| | | | | | | |
|---|---|---|---|---|---|---|
| *ANG* | Increased proliferative capacity of hematopoietic stem/progenitor cells and decreased proliferative capacity of myeloid-restricted progenitor cells and develop leukopenia | Amyotrophic lateral sclerosis 9 | Missense (mainly) | LOF | Amyotrophic lateral sclerosis was an adult-onset, rapidly progressive, and ultimately fatal neurodegenerative disorder, which included weakness, muscular atrophy, and spasticity that may evolve to paralysis. | 17900154 |
| *ANO10* | Fail to exhibit calcium-activated chloride ion secretion in the jejunum | Spinocerebellar ataxia, autosomal recessive 10 | Missense & destructive | LOF | The disorder was characterized by slowly progressive spastic ataxia variably associated with motor neuron involvement, epilepsy, and cognitive decline. Age at onset ranged between 27 and 53 years. | 30515630 |
| *AP5Z1* | Abnormal head morphology | Spastic paraplegia 48, autosomal recessive | Missense & destructive | LOF | The disorder was characterized by spasticity of the lower limbs resulting in gait difficulties. Most patients were onset in mid- or late-adulthood. | 26085577 |
| *DNAJB2* | Altered total body fat amount | Spinal muscular atrophy, distal, autosomal recessive, 5 | Missense & destructive | LOF | The disorder was characterized by adult onset of slowly progressive distal muscle weakness and atrophy resulting in gait impairment and loss of reflexes due to impaired function of motor nerves. | 22522442 |
| *DRAM2* | Abnormal coat/hair pigmentation | Cone-rod dystrophy 21 | Missense & destructive | LOF | The disorder was adult-onset, characterized by retinal pigment deposits visible on fundus examination, predominantly in the macular region, and initial loss of cone photoreceptors followed by rod degeneration. | 25983245 |
| *EIF2B5* | Decreased circulating glucose level, heart weight, mean corpuscular hemoglobin concentration, and mean corpuscular volume | Leukoencephalopathy with vanishing white matter | Missense & destructive | LOF | Vanishing white matter leukodystrophy was a neurologic disorder characterized by variable neurologic features. Individuals with *EIF2B5* were adult onset. | 21484434 |
| *MAK* | Slight reductions in litter size and sperm motility | Retinitis pigmentosa 62 | Missense & destructive | LOF | Among the 8 patients reported, the age at diagnosis in 3 patients were in the third decade of life, whereas the other 5 patients were diagnosed in the fourth through sixth decades of life. | 21835304 |
| *MUTYH* | Decreased susceptibility to neuronal excitotoxicity | Adenomas, multiple colorectal | Missense & destructive | LOF | The disorder was characterized by adult-onset of multiple colorectal adenomas and adenomatous polyposis. Affected individuals had a significantly increased risk of colorectal cancer. | 12606733 |
| *RTN2* | Impaired glucose uptake in skeletal muscle | Spastic paraplegia 12, autosomal dominant | Missense & destructive | LOF | The disorder was characterized by lower limb spasticity and hyperreflexia, resulting in walking difficulties. The age at onset was variable and can range from childhood to adulthood. | 22232211 |

| | | | | | | |
|---|---|---|---|---|---|---|
| **4. Unexplainable (n=5)** | | | | | | |
| *AP4M1* | Decreased circulating serum albumin level, decreased total body fat amount, and improved glucose tolerance | Spastic paraplegia 50, autosomal recessive | Destructive (mainly) | LOF | The disorder was characterized by neonatal hypotonia that progressed to hypertonia and spasticity and severe mental retardation with poor or absent speech development. | 19559397 |
| *DNAJC19* | Decreased body weight, decreased lean body mass, and increased circulating glucose level | 3-methylglutaconic aciduria, type V | Destructive | LOF | The disorder was characterized by the onset of dilated or noncompaction cardiomyopathy in infancy or early childhood. | 22797137 |
| *FAM126A* | Decreased expression of TTC7A and EFR3A. | Leukodystrophy, hypomyelinating, 5 | Missense & destructive | LOF | The disorder was characterized by hypomyelination of the central and peripheral nervous system, progressive neurological impairment and congenital cataract. | 26571211 |
| *FAR1* | Decreased B cell number | Peroxisomal fatty acyl-CoA reductase 1 disorder | Missense & destructive | LOF | The disorder was characterized by onset in infancy of severely delayed psychomotor development, growth retardation with microcephaly, and seizures. | 25439727 |
| *TMEM165* | Increased mean corpuscular volume and abnormal iris morphology | Congenital disorder of glycosylation, type IIk | Missense & destructive | LOF | Affected individuals showed psychomotor retardation and growth retardation, and most had short stature. Other features included dysmorphism, hypotonia, eye abnormalities, acquired microcephaly, hepatomegaly, and skeletal dysplasia. | 22683087 |

Abbreviations: DNE, dominant-negative effect; GOF, gain of function; LOF, loss of function.

**Table S2. The genes of quantitatively defined genetic dependent quantity (GDQ) and genetic dependent range (GDR)**

| Gene | Quantitative correlation (evidence from residual activity, unless otherwise specified) | GDQ | GDR | Reference (PMID) |
|---|---|---|---|---|
| *SCN1A* | 50.0%-75%–a variety of epilepsies, complete penetrance<br>75%-87.5%–febrile seizure or mild epilepsy, incomplete penetrance<br>83%–highest level identified in patients<br>>87.5%–asymptomatic<br>(mosaic evidence) | 83% | 50%-83% | 25754450 |
| *LDLR* | <2%–severe familial hypercholesterolaemia<br>>2%–mild familial hypercholesterolaemia<br>80%-normal | 80% | 2%-80% | 28169869 |
| *FLNB* | 50% (heterozygous)–classic atelosteogenesis, type I<br>80%–mild limb deformities and corporal asymmetry<br>(mosaic evidence) | 80% | 50%-80% | 30231296 |
| *KCNQ2* | 50% (missense with dominant negative effect)–developmental and epileptic encephalopathy<br>75%–benign familial neonatal epilepsy | 75% | 50%-75% | 20437616 |
| *SLC2A1* | 50%–classic glucose transporter type 1 deficiency syndrome<br>65%–paroxysmal events such as seizures, exercise-induced dyskinesias, and intermittent ataxia<br>>75%–few or no clinical symptoms | 75% | 50%-75% | 20687207 |
| *CREBBP* | 70%–typical Rubinstein-Taybi syndrome<br>(mosaic evidence) | >70% | - | 17855048 |
| *SLC16A1* | 42.1%-59.3%–symptomatic deficiency in lactate transport | ~60% | - | 10590411 |
| *ZSWIM6* | 50% (heterozygous)–neurodevelopmental disorder with movement abnormalities, abnormal gait, and autistic features<br>60%–mild acromelic frontonasal dysostosis<br>(mosaic evidence) | 60% | 50%-60% | 25105228 |
| *COL1A1* | 30%-50%–mild osteogenesis imperfecta<br>57%-89%–normal<br>(mosaic evidence) | 50% | 30%-50% | 12538651,<br>8799376 |
| *COL7A1* | <30%–generalized dystrophic epidermolysis bullosa<br>~50% (heterozygous with dominant-negative effect)–localized dystrophic epidermolysis bullosa<br>~75%–normal<br>(mosaic evidence) | 50% | 30%-50% | 9668111,<br>24317394,<br>25113066 |
| *TUBB4A* | 48%–hypomyelination with atrophy of the basal ganglia and cerebellum<br>50% (heterozygous with dominant negative effect)–hypomyelinating leukodystrophy<br>(mosaic evidence) | 50% | - | 23582646 |
| *ATP7A* | 0-15%–Menkes disease<br>20-35%–occipital horn syndrome<br>~50% (heterozygous females)–usually unaffected, occasionally Menkes disease | 50% | 0-50% | 20301586,<br>20497190 |
| *CHST3* | 29%–Omani-type spondyloepiphyseal dysplasia with cardiac involvement<br>~50% (heterozygous)–usually unaffected, occasionally lumbar disc degeneration | 50% | 18%-50% | 24216480,<br>19320654 |
| *CPA6* | 40% (homozygous)–febrile seizures, familial, 11<br>50% (heterozygous)–epilepsy, familial temporal lobe, 5 | 50% | - | 21922598 |

| | | | | |
|---|---|---|---|---|
| *CPOX* | 49%–hereditary coproporphyria<br>50% (heterozygous)–hereditary coproporphyria or asymptomatic<br>58%–asymptomatic | 50% | - | 12227458 |
| *MTHFR* | <36%–methylenetetrahydrofolate reductase deficiency<br>50% (heterozygous)–susceptibility | 50% | - | 12673793 |
| *NAGLU* | 0%-3%–severe mucopolysaccharidosis type IIIB<br>50% (heterozygous)–mild Charcot-Marie-Tooth disease, axonal, type 2V | 50% | 0-50% | 25818867 |
| *PRPS1* | 4%-10% (in males) –severe Charcot-Marie-Tooth disease-5<br>18.3%-47.7% (in carrier females)–mild Charcot-Marie-Tooth disease-5<br>25%-35%–X-linked deafness-1 | 50% | 4%-50% | 25182139 |
| *RRM2B* | 1%-2%–severe mitochondrial DNA depletion syndrome<br>12%–mitochondrial neurogastrointestinal encephalopathy syndrome<br>50%–progressive external ophthalmoplegia with mitochondrial DNA deletions | ~50% | 1%-50% | 17486094,<br>9667227,<br>21646632 |
| *SLC16A2* | 20%-37%–Allan-Herndon-Dudley syndrome<br>50% (heterozygous females)–mild thyroid phenotype without neurologic defects | 50% | 20%-50% | 18187543 |
| *SLC25A4* | <1%–mitochondrial DNA depletion syndrome 12B<br>~50% (heterozygous with dominant negative effect)–progressive external ophthalmoplegia with mitochondrial DNA deletions | 50% | 0-50% | 17486094,<br>19664747 |
| *SLC26A2* | 0–very severe achondrogenesis type IB<br>40%–severe diastrophic dysplasia<br>50%–mild multiple epiphyseal dysplasia-4 | 50% | 0-50% | 15294877 |
| *STAR* | 3%-10%–severe congenital adrenal hyperplasia<br>50.0%–mild congenital adrenal hyperplasia | 50% | 3%-50% | 20444910 |
| *UROD* | 5%-20%–hepatoerythropoietic porphyria<br>~50%–susceptible to porphyria cutanea tarda | 50% | 5%-50% | 24175354,<br>17360334 |
| *NDUFS1* | 45%–mitochondrial complex I deficiency nuclear type 5<br>50% (heterozygous)–asymptomatic | 45% (<50%) | - | 21203893 |
| *TK2* | 14%-45% (in muscle)–mitochondrial DNA depletion syndrome<br>50% (heterozygous)–asymptomatic | ~45% | - | 11687801 |
| *C3* | 0.1%–C3 deficiency<br>>43.1%–normal | <43% | - | 15781264 |
| *GPI* | 0%-42%–GPI deficiency<br>50% (heterozygous)–asymptomatic | 42% | 0-42% | 19064002 |
| *INPP5K* | 27%~42%–congenital muscular dystrophy with cataracts and intellectual disability<br>40%-60% (heterozygous)–asymptomatic | 42% | - | 28190459 |
| *CYP11B1* | 0-5%–classic 11beta-hydroxylase deficiency (androgen excess, virilization, and hypertension)<br>9-40%–non-classic 11beta-hydroxylase deficiency (without signs of virilisation)<br>50% (heterozygous)–asymptomatic | 40% | 0~40% | 29697234 |
| *NUBPL* | 40%–mitochondrial complex I deficiency nuclear type 21<br>50% (heterozygous)–asymptomatic | 40% (<50%) | - | 20818383 |
| *TTC19* | 8%-39%–mitochondrial complex III deficiency nuclear type<br>50% (heterozygous)–asymptomatic | 39% (<50%) | - | 23532514 |

| | | | | |
|---|---|---|---|---|
| *DHCR7* | 0-8.2%–Smith-Lemli-Opitz syndrome<br>14.5%-37%–very mild Smith-Lemli-Opitz syndrome<br>50% (heterozygous)–asymptomatic | 37% | 0-37% | 20301322, 31088393 |
| *ASPA* | <5%–most severe Canavan disease<br>20%-35%–milder Canavan disease<br>50% (heterozygous)–asymptomatic | 35% | 5%-35% | 22850825 |
| *ABCA4* | 0–severe Stargardt disease<br>34.3%–moderate Stargardt disease<br>(splicing evidence) | 34.3% | 0-34.3% | 29162642 |
| *DLD* | 0%-33%–dihydrolipoamide dehydrogenase deficiency<br>42%-44%–asymptomatic | 33% | 0-33% | 16770810, 8968745 |
| *QARS1* | 33%–severe growth deficiency, microcephaly, intellectual disability, and facial features<br>50% (heterozygous)–asymptomatic | ~33% (<50%) | - | 28620870 |
| *ASAH1* | <5%–classic Farber disease<br><10%–Farber disease<br>~32%–spinal muscular atrophy with progressive myoclonic epilepsy | 32% | 5%-32% | 22703880 |
| *GLDC* | 0–severe nonketotic hyperglycinemia<br>6.1%-32%–mild nonketotic hyperglycinemia | 32% | 0-32% | 15236413, 15824356 |
| *BTD* | <10%–profound biotinidase deficiency<br>10-30%–partial biotinidase deficiency (often asymptomatic, unless when stressed)<br>45%-50%–asymptomatic | 30% | 10%~30% | 20301497 |
| *GAA* | <3%–infants with classic Pompe's disease<br>3%-30%–children and adults with less severe Pompe's disease<br>~50%–unaffected carriers | 30% | 0-30% | 18929906, 24399863 |
| *KCNJ10* | <30%–SESAME syndrome<br>30%–asymptomatic | 30% | - | 20807765 |
| *NDUFA1* | 20%–severe mitochondrial complex I deficiency, nuclear type 12<br>30%–mild mitochondrial complex I deficiency, nuclear type 12<br>50% (heterozygous females)–asymptomatic | 30% | 20%-30% | 17262856 |
| *PHKA2* | 2%-16.9%–glycogen storage disease type IXa1<br>30%–mild glycogen storage disease type IXa1 | 30% | 2%-30% | 7711737 |
| *UGT1A1* | 0–Crigler-Najjar syndrome type 1<br><20%–Crigler-Najjar syndrome type 2<br>30%–Gilbert syndrome (often asymptotic) | 30% | 0-30% | 8528206, 11906189, 7989595 |
| *ACADVL* | <10%–deficiency of very long-chain acyl-CoA dehydrogenase<br>10%-30%–risk of symptoms in situations of increased energy demand and during illnesses<br>>30.0%–asymptomatic | 30% | 10%-30% | 21932095 |
| *BCKDHA* | <2%–classic maple syrup urine disease<br>3%-15%–intermediate maple syrup urine disease<br>17%-30%–very mild clinical course of maple syrup urine disease | 30% | 2%-30% | 22727569, 16786533, 17922217 |
| *ADSL* | 30%–adenylosuccinase deficiency<br>50% (heterozygous)–asymptomatic | ~30% (<50%) | - | 15471876 |

| | | | | |
|---|---|---|---|---|
| *AK1* | 30%–severe congenital adenylate kinase deficiency<br>50% (heterozygous)–asymptomatic | ~30% (<50%) | - | 30918013 |
| *MLYCD* | 30.3%–malonyl-CoA decarboxylase deficiency<br>50% (heterozygous)–asymptomatic | ~30% (<50%) | - | 12955715 |
| *MRPS22* | 8%-30%–combined oxidative phosphorylation deficiency 5<br>50% (heterozygous)–asymptomatic | ~30% (<50%) | - | 17873122 |
| *PDHX* | 30%–pyruvate dehydrogenase complex deficiency<br>50% (heterozygous)–asymptomatic | ~30% (<50%) | - | 17152059 |
| *PLA2G6* | 30%–*PLA2G6*-associated neurodegeneration<br>50% (heterozygous)–asymptomatic | ~30% (<50%) | - | 21700586 |
| *SDHD* | 30% (skeletal muscle)–mitochondrial complex II deficiency<br>50% (heterozygous)–asymptomatic | ~30% (<50%) | - | 26008905 |
| *CTNS* | 0-5%–infantile cystinosis<br>5-29%–mild ocular cystinosis/nephropathic cystinosis<br>50% (heterozygous)–asymptomatic | 29% | 0-29% | 11505338,<br>15128704,<br>12110740 |
| *SLC25A20* | <5%–severe carnitine-acylcarnitine translocase deficiency<br>27%–mild carnitine-acylcarnitine translocase deficiency | 27% | 5%-27% | 11162577 |
| *NDUFA11* | 13%-27%–mitochondrial complex I deficiency nuclear type 14<br>50% (heterozygous)–asymptomatic | ~27% (<50%) | - | 18306244 |
| *CLPB* | 26%–3-methylglutaconic aciduria, with cataracts, neurologic involvement and neutropenia<br>50% (heterozygous) –asymptomatic | 26% (<50%) | - | 25597510 |
| *CPT2* | 2%–perinatal onset carnitine palmitoyltransferase II deficiency<br>5%–lethal neonatal form of CPT II deficiency<br>15%-26%–adult onset CPT II deficiency<br>35%-65%–asymptomatic (carrier) | 26% | 2%-26% | 18550408,<br>15622536,<br>14615409,<br>1528846 |
| *CPS1* | 0%-39%–carbamoyl phosphate synthetase I deficiency<br>10%-25%–less severe hyperammonemia<br>50%–normal | 25% | 0-25% | 17310273,<br>12955727 |
| *GALT* | 0%-15%–classic galactosemia<br>25%–Duarte variant galactosemia<br>50% (heterozygous)–asymptomatic | 25% | 0-25% | 25473725,<br>20301691 |
| *ITGB3* | 5%–most severe type I Glanzmann thrombasthenia<br>10-25%–moderate type II Glanzmann thrombasthenia<br>50% (heterozygous)–asymptomatic | 25% | 5%-25% | 31029159 |
| *PTF1A* | 0–pancreatic and cerebellar agenesis<br>25%–isolated pancreatic agenesis | 25% | 0-25% | 27284104 |
| *PMM2* | 0–severe congenital disorder of glycosylation, type Ia<br>>25%–mild or very mild phenotype<br>50% (heterozygous)–asymptomatic | 25% | 0-25% | 30740725,<br>11156536 |
| *SGSH* | <3.0%–severe children onset mucopolysaccharidosis type IIIA<br>3.0-11.0%–mild children onset mucopolysaccharidosis type IIIA<br>25.0%–adults onset mucopolysaccharidosis type IIIA | 25% | 3%-25% | 21671382 |

| | | | | |
|---|---|---|---|---|
| *TYR* | 0–oculocutaneous albinism type IA<br>12%-13%–oculocutaneous albinism type IB<br>25%–unaffected or oculocutaneous albinism type IB | 25% | 0~25% | 19208379,<br>18326704,<br>1899321 |
| *ACADS* | 25%–short-chain acyl-CoA dehydrogenase deficiency<br>40%–asymptomatic | ~25% (<40%) | - | 9499414,<br>20376488 |
| *ACAT1* | 25%–mitochondrial acetoacetyl-CoA thiolase deficiency<br>50% (heterozygous)–asymptomatic | ~25% (<50%) | - | 17236799 |
| *FECH* | 4%-25%–erythropoietic protoporphyria<br>50% (heterozygous)–asymptomatic | ~25% (<50%) | - | 17875872,<br>1184741,<br>7356682 |
| *PC* | 2-25%–pyruvate carboxylase deficiency<br>50% (heterozygous)–asymptomatic | ~25% (<50%) | - | 9585002 |
| *CYP27B1* | 0-22%–vitamin D-dependent rickets<br>50% (heterozygous)–asymptomatic | ~22% (<50%) | - | 12050193 |
| *DPAGT1* | 10%-22%–congenital disorder of glycosylation type Ij<br>50% (heterozygous)–asymptomatic | ~22% (<50%) | - | 22492991,<br>12872255 |
| *SMPD1* | <5%–Niemann-Pick disease type A<br>21.5%–later onset and milder Niemann-Pick disease type B | 21.5% | 5%-21.5% | 19405096 |
| *POMGNT1* | <1%–uscular dystrophy-dystroglycanopathy (congenital with brain and eye anomalies)<br>21%–non-syndromic retinitis pigmentosa | 21% | 0-21% | 11709191,<br>27391550 |
| *ARG1* | 0%-20.8%–hyperargininemia<br>50% (heterozygous)–asymptomatic | ~21% (<50%) | - | 22959135 |
| *CPT1A* | <10%–classic carnitine palmitoyltransferase 1 deficiency<br>20%–mild symptoms (muscle cramps that recovered spontaneously within 1 day)<br>~50% (heterozygous)–asymptotic | 20% | 0~20% | 11441142,<br>20301700 |
| *CYP21A2* | 0-11%–classical form of congenital adrenal hyperplasia<br>20.0%–non-classical congenital adrenal hyperplasia<br>50% (heterozygous)–asymptomatic | 20% | 0~20% | 20301350 |
| *DHCR24* | <1%–severe desmosterolosis<br>20%–mild desmosterolosis<br>50% (heterozygous)–asymptomatic | 20% | 0-20% | 11519011,<br>21671375 |
| *ADK* | 20%–hypermethioninemia due to adenosine kinase deficiency<br>50% (heterozygous)–asymptomatic | ~20% (<50%) | - | 21963049 |
| *CA5A* | 20%–carbonic anhydrase VA deficiency<br>50% (heterozygous)–asymptomatic | ~20% (<50%) | - | 24530203 |
| *DPM1* | 8%-20%–congenital disorder of glycosylation type Ie<br>50% (heterozygous)–asymptomatic | ~20% (<50%) | - | 16641202,<br>23856421 |
| *SLC25A15* | 4%-19%–hyperornithinemia-hyperammonemia-homocitrullinuria syndrome<br>50% (heterozygous)–asymptomatic | >19% (<50%) | - | 19242930 |
| *CFTR* | 6.9%–classic cystic fibrosis<br>11.2%–milder cystic fibrosis<br>18.9%–isolated congenital absence of the vas deferens | 19% | 7%-19% | 27738188 |
| *CFTR* | 13%–isolated congenital absence of the vas deferens | 13% | - | 15705292 |

| | | | | |
|---|---|---|---|---|
| | 19%–asymptomatic<br>(splicing evidence) | | | |
| *COQ9* | 18%–primary Coenzyme Q10 deficiency<br>50% (heterozygous)–asymptomatic | 18% (<50%) | - | 20495179 |
| *PGK1* | 1.6%-17.5%–phosphoglycerate kinase-1 deficiency<br>50% (heterozygous females)–asymptomatic | ~18% (<50%) | - | 19157875 |
| *PCCA* | 3.3%–late-onset, relatively mild form of propionic academia<br>16%–slight psychomotor delay<br>(splicing evidence) | 16% | 3.3%-16% | 15235904 |
| *SUMF1* | 0–neonatal multiple sulfatase deficiency<br>1.6%–late infantile severe multiple sulfatase deficiency<br>16%–late infantile and mild multiple sulfatase deficiency | 16% | 0-16% | 21224894 |
| *TYMP* | <5%–mitochondrial DNA depletion syndrome-1<br>9%-16%–ate onset mitochondrial DNA depletion syndrome-1<br>26%-35%–asymptomatic | 16% | 5%-16% | 9924029,<br>15975738,<br>16178026 |
| *GINS1* | 3%-16%–immunodeficiency<br>50% (heterozygous)–asymptomatic | ~16% (<50%) | - | 28414293 |
| *MUTYH* | 5%–familial adenomatous polyposis<br>15%–mild familial adenomatous polyposis<br>50% (heterozygous)–asymptomatic | 15% | 5%-15% | 20848659,<br>19032956 |
| *SLC25A12* | 15%–developmental and epileptic encephalopathy<br>50% (heterozygous) –asymptomatic | ~15% (<50%) | - | 24515575 |
| *HADHB* | 0–severe trifunctional protein deficiency<br>5%-14%–mild trifunctional protein deficiency | 14% | 0-14% | 19699128 |
| *MT-ND3* | 1%–lethal infantile mitochondrial disease<br>14%–leigh syndrome<br>50%-74%–asymptomatic | ~14% (<50%) | - | 11456298,<br>15372108,<br>17152068 |
| *D2HGDH* | 0–severe D-2-hydroxyglutaric<br>13%–mild form D-2-hydroxyglutaric<br>50% (heterozygous)–asymptomatic | 13% | 0-13% | 16037974,<br>30908763 |
| *GALNS* | 0–severe mucopolysaccharidosis IVA<br>1.3%-13.3%–mild mucopolysaccharidosis IVA | 13% | 0-13% | 10814710 |
| *GLB1* | 0–infantile GM1-gangliosidosis, type I<br>~1%-5%–late infantile GM1-gangliosidosis, type II<br>~3%-10%–juvenile GM1-gangliosidosis, type II<br>5%-10%– chronic/adult GM1-gangliosidosis, type III<br>2%-12%–mucopolysaccharidosis, type IVB | 12% | 0-12% | 24156116,<br>12644936 |
| *GLUL* | 12%–congenital glutamine deficiency<br>50% (heterozygous)–asymptomatic | ~12% (<50%) | - | 16267323 |
| *CTSD* | 0–severe neuronal ceroid lipofuscinosis<br>11%–less severe neuronal ceroid lipofuscinosis<br>50% (heterozygous)–asymptomatic | 11% | 0-11% | 16670177,<br>25298308 |
| *HEXA* | 0.1%–Tay-Sachs disease<br>0.5%–late infantile GM2-gangliosidosis | ~11% | 0-11% | 6614006,<br>20301397 |

| | | | | |
|---|---|---|---|---|
| | 2%-4%–adult GM2-gangliosidosis<br>11%-20%–healthy persons with low hexosaminidase | | | |
| *NDUFA12* | 11%–mitochondrial complex I deficiency nuclear type 23<br>50% (heterozygous)–asymptomatic | ~11% (<50%) | - | 21617257 |
| *ADAMTS13* | <10.0%–thrombotic thrombocytopenic purpura<br>>10%–thrombotic thrombocytopenic purpura (recurrence risk of 4%)<br>50% (heterozygous)–asymptomatic | 10% | - | 23415418, 16420561 |
| *ALAD* | 10%–acute hepatic porphyria<br>12%–asymptomatic | 10% | - | 15303011, 10519994 |
| *ARSA* | 5.0%-10.0%–metachromatic leukodystrophy<br>10.0%-30.0%–pseudodeficiency (very low levels of arylsulfatase activity in healthy individuals)<br>>30.0%–normal | 10% | 0-10% | 1348043, 24411407 |
| *DHFR* | 0–severe dihydrofolate reductase deficiency<br>~10%–mild dihydrofolate reductase deficiency | 10% | 0-10% | 21310276, 21310277 |
| *GNPTAB* | <1%–severe mucolipidosis II alpha/beta<br>1-10%–mild mucolipidosis III alpha/beta | 10% | 0-10% | 19617216, 20301728 |
| *ITPA* | 0.3%-5.3%–developmental and epileptic encephalopathy 35<br>10%–asymptomatic | 10% | 0-10% | 26224535, 12384777 |
| *SLC25A13* | 10%–adult-onset type 2 citrullinemia | 10% | - | 18367750 |
| *UROS* | <1%–non-immune hydrops fetalis<br>~10%–congenital erythropoietic porphyria<br>50% (heterozygous)–asymptomatic | 10% | 0-10% | 24027798 |
| *ALG1* | <10%–congenital disorder of glycosylation, type Ik<br>50% (heterozygous)–asymptomatic | ~10% (<50%) | - | 20679665 |
| *AP1S1* | <10% (mRNA)–MEDNIK syndrome<br>40%-75% (mRNA)–asymptomatic | ~10% (<40%) | - | 19057675 |
| *CAT* | <10%–acatalasemia<br>50% (heterozygous)–asymptomatic | ~10% (<50%) | - | 24522161 |
| *MAN2B1* | 5%-10%–alpha-mannosidosis<br>50% (heterozygous)–asymptomatic | ~10% (<50%) | - | 20301570 |
| *PROS1* | <10%–thrombophilia due to protein S deficiency<br>50% (heterozygous)–asymptomatic | ~10% (<50%) | - | 20484936 |
| *PSPH* | <10%–phosphoserine phosphatase deficiency<br>50% (heterozygous)–asymptomatic | ~10% (<50%) | - | 25080166 |
| *SCARB2* | 10%–action myoclonus-renal failure syndrome<br>50% (heterozygous)–asymptomatic | ~10% (<50%) | - | 18424452 |
| *ALDOA* | 4%-9%–glycogen storage disease XII<br>50% (heterozygous)–asymptomatic | ~9% (<50%) | - | 8598869, 25392908 |
| *HPRT1* | <1.5%–Lesch-Nyhan disease<br>1.5%-8%–HGprt-related neurological dysfunction<br>>8%–mild HGprt-related hyperuricemia | ~8% | 1.5%-8% | 22157001 |
| *MVK* | 0–severe mevalonic aciduria<br>1%-8%–mild hyperimmunoglobulinemia D with periodic fever syndrome | 8% | 0-8% | 16835861 |

| | | | | |
|---|---|---|---|---|
| *PEPD* | 0–severe prolidase deficiency<br>7.4%–mild prolidase deficiency | 7.40% | 0-7.4% | 16470701,<br>8900231 |
| *ARSB* | 2%–severe mucopolysaccharidosis type VI<br>7%–mild mucopolysaccharidosis type VI<br>50% (heterozygous)–asymptomatic | 7% | 0-7% | 1550123 |
| *NDUFA10* | 7%–mitochondrial complex I deficiency, nuclear type 22<br>50% (heterozygous)–asymptomatic | >7% (<50%) | - | 21150889 |
| *ATR* | 6%–Seckel syndrome<br>47%–asymptomatic<br>(splicing evidence) | >6% (<47%) | - | 27639833 |
| *NAGA* | 0%-5.9%–Schindler disease<br>6%-21%–asymptomatic | 6% | 0-6% | 31468281,<br>11313741 |
| *PGAM2* | 2.1%–glycogen storage disease X<br>6%–mild glycogen storage disease X | 6% | 2.1%-6% | 8447317 |
| *TPP1* | 0–severe neuronal ceroid lipofuscinosis with greatly shortened lifespan<br>3%–delayed disease onset and longer survival<br>6%–dramatically attenuated disease | 6% | 0-6% | 18343701 |
| *GUSB* | 2%-5% (homozygous)–mucopolysaccharidosis VII<br>40%-60% (heterozygous)–asymptomatic | ~5% (<50%) | - | 1702266,<br>9987917 |
| *DOLK* | 2%-4%–congenital disorder of glycosylation type Im<br>50% (heterozygous)–asymptomatic | ~4% (<50%) | - | 17273964 |
| *DDC* | <1%–aromatic L-amino acid decarboxylasedeficiency<br>2.7%-3.7%–AADC deficiency (in drosophila)<br>40%–asymptomatic | >3.7%<br>(<40%) | - | 14991824 |
| *F7* | 3%–factor VII deficiency<br>50% (heterozygous)–asymptomatic<br>(splicing evidence) | >3% (<50%) | - | 31273093 |
| *PYGM* | 0–typical glycogen storage disease type<br>1%-2.5%–mild glycogen storage disease | 2.50% | 0-2.5% | 19433441 |
| *IDS* | 0–severe mucopolysaccharidosis II<br>0.2%-2.4%–mild mucopolysaccharidosis I | 2.40% | 0-2.4% | 17091340 |
| *GLA* | <1%–classic Fabry disease<br>>1%–late-onset atypical Fabry disease, cardiac variant | ~1% | 0-1% | 20301469 |
| *ADA* | <0.05%–severe combined immunodeficiency<br>0.06%-0.6%–late onset combined immunodeficiency<br>1%-1.5%–effective immune function | 1% | 0-1% | 14499267 |

Note: the presented GDRs were based on clinically observed phenotypes. In genes of vital in GDN, the severe extreme of phenotype spectrum were embryonic or early death, which would be less clinically diagnosed.
